# Methamphetamine and α-pyrrolidinopentiophenone (α-PVP) Intravenous Self-Administration in Female and Male Rats

**DOI:** 10.1101/2025.09.12.675933

**Authors:** Arnold Gutierrez, Yanabel Grant, Sophia A. Vandewater, Michael A. Taffe

## Abstract

**Background:** Stimulant drug users vary in their substance of choice and may, in some cases, switch up their preferred substance based on availability, cost or other factors. Poly-substance use is rarely assessed in rodent models of drug seeking and this study determined if training drug alters the apparent reinforcing properties of methamphetamine (MA) and α-pyrrolidinopentiophenone (α-PVP).

**Methods:** Female and male Wistar rats (N=8 per group) were trained in the intravenous self-administration (IVSA) of α-PVP or MA. The impact of dose substitution (0.0125, 0.0250, 0.100, 0.300 mg/kg/infusion) for each training drug was then assessed in all groups under FR and Progressive Ratio schedules of reinforcement.

**Results:** Male and female rats obtained similar numbers of infusions of MA (0.05 mg/kg/infusion) and of α-PVP (0.05 mg/kg/infusion) during acquisition, however more infusions of α-PVP than of MA were obtained by each sex. Mean lever discrimination ratios exceeded 80% on the drug-associated lever within 5 training sessions for α-PVP groups but were not consistently at this level for either MA group. Drug potency was similar across groups but was less effective in the MA-trained males.

**Conclusions:** Interpretations of sex differences in the acquisition of drug IVSA require caution when dose is not varied across or within group. This study also further confirms that the apparent efficacy of a drug as a reinforcer depends at least partially on the behavioral antecedents, including the identity of the drug used for initial IVSA acquisition.

## 1. Introduction

Nonmedical use of psychostimulant drugs, including methamphetamine (**MA**), cocaine and synthetic cathinones, has led to substantive individual and public health concerns worldwide (Webb et al., 2024). In the United States, the latest available data from the National Survey on Drug Use and Health reports 2.7 million people used MA in the past year and 1.8 million people had a MA use disorder (SAMHSA, 2023). This rate was higher than the 1.4 million with a cocaine disorder out of the 5.3 million who used cocaine in the past year (SAMHSA, 2023). Only 476 thousand received inpatient care for a MA use disorder and only 708,000 received outpatient care. Relapse rates are high for those who do enter treatment, for example 61% of individuals who exited an inpatient treatment program returned to MA use within one year, and only 13% remained abstinent for over 5 years (Brecht and Herbeck, 2014). MA was the most commonly reported substance in the 2019 National Forensic Laboratory Information System (NFLIS) data set, accounting for 27% of all reported items, outpacing cocaine (14%). Relatedly, MA -involved deaths in the US increased five-fold from 2011 to 2018 (Han et al., 2021).

The emergence of a variety of synthetic cathinones over the past 15 years presented a moving target, as new analogs waxed and waned in recreational availability (DEA, 2014, 2019, 2020; Mohr et al., 2018) in a presumed effort to evade legal control. Compounds such as 3,4-methylenedioxypyrovalerone (**MDPV**; “bath salts”, “monkey dust”) and α-pyrrolidinopentiophenone (or α-pyrrolidinovalerophenone, α-PVP; “flakka”) act as restricted monoamine transporter inhibitors with activity primarily at the dopamine and norepinephrine transporters (Baumann et al., 2013; Kolanos et al., 2015; Simmler et al., 2013). Unsurprisingly due to this neuropharmacology, MDPV, α-pyrrolidinopentiophenone (**α-PVP**) and associated close analogs of these compounds have substantial propensity for compulsive use, binging and psychotomimetic behavior. The National Forensic Laboratory Information System 2019 annual report and the December 2020 and December 2024 updates report α-PVP analogs α-PHP and α-PiHP continued to be popular (DEA, 2020, 2021, 2024) and there are continued arrests for MDPV possession and trafficking (CBS17, 2020; Fox12Staff, 2019; Holsman, 2021; Nash, 2024). Correspondingly, drugs in this class are highly effective reinforcers in laboratory models of drug misuse. MDVP and α-PVP are approximately equally effective and potent in self-administration models (Aarde et al., 2015a; Collins et al., 2019; Gannon et al., 2017; Javadi-Paydar et al., 2018a), and many close analogs of each compound differ very little in similar pre-clinical investigations (Gannon et al., 2018a; Gannon et al., 2018b; Huskinson et al., 2017).

Problematic use of a given psychostimulant drug occurs in some individuals with prior or concurrent patterns of use of other psychostimulants including MA and cocaine, but also synthetic cathinones (Albert et al., 2022; Barton and Wang, 2023; Bender et al., 2022; Enabi et al., 2024; Lin et al., 2023; Mohr et al., 2018). Users of multiple psychostimulants, including cathinones and MA, may exhibit poorer health and greater complications from acute intoxication compared with MA-only users (Lin et al., 2023). Evaluation of the impact of exposure to more than one psychostimulant is rare in animal models, but some prior work suggests that the efficacy of one drug as a reinforcer in intravenous self-administration (**IVSA**) may be altered by prior exposure to a different psychostimulant (Creehan et al., 2015; Khom et al., 2021; Seaman et al., 2022). This suggests that risks of a given stimulant may depend on the antecedent drug use history of the individual as much, or more so, than the inherent properties of the drug in question. Thus, one major goal of this study was to determine if initial IVSA training with MA alters the IVSA of α-PVP and *vice versa*.

A review by Anker and Carroll summarizes evidence from rodent self-administration studies to conclude that female animals are at higher risk for abuse-related profiles of stimulant drug self-administration (Anker and Carroll, 2011). They cite evidence that female rats acquire MA (0.02 mg/kg/infusion) self-administration faster, as defined by a set total number of drug infusions (Roth and Carroll, 2004), and obtain more infusions across a range of doses (0.01-0.08 mg/kg/infusion) post-acquisition in a Progressive Ratio procedure. They also cite additional evidence that female rats escalate their drug intake to a greater degree in extended access paradigms. Broader availability of self-administration data that examines the contribution of rat sex in more recent years, due in part to the Sex as a Biological Variable (SABV) initiative promulgated by the U.S. National Institutes of Health (Clayton and Collins, 2014; Shansky and Murphy, 2021), suggests a less unified conclusion. For example, male Sprague-Dawley (SD) rats obtained more infusions of MA (0.1 mg/kg/infusion) than did female SD rats (Funke et al., 2023) at the end of a 10 day acquisition (in 6 h access sessions in the dark cycle). Male Long-Evans rats obtained more infusions of MA (0.1 mg/kg/infusion) than female rats did when trained in two daily 3 h sessions (Miller et al., 2022), as did male Wistar rats trained in two 2 h daily sessions for MA (0.05 mg/kg/infusion) IVSA in the light cycle (Zlebnik et al., 2021). Comparison of prior findings can be complicated due to methodological variations. For example, Long-Evans female rats obtained more MA deliveries than did male rats in a 6 h IVSA access procedure in the dark (Reichel et al., 2012), but the authors used a 20 μg bolus infusion in males and a 17.5 μg meth bolus in females to account for body size; females therefore received 87.5% of the male dose with reported entry weights that were 66.7% of the male weight. Despite the fact that most IVSA sex-comparisons adjust the per-infusion dose by body weight throughout the study, it could still be the case that this approximation fails to equalize brain drug levels. If this method consistently provided females with lower brain drug levels for a given number of infusions, this might explain the apparent sex difference. This concern is countered in some studies with the inclusion of dose-substitution procedures in which an effective potency difference can be distinguished from apparent efficacy difference across the dose range. Although available IVSA data are more limited for α-PVP or MDPV, there were no differences in male and female self-administration of α-PVP (0.1 mg/kg/infusion) in either 1 h or 6 h daily sessions (Marusich et al., 2021) nor any sex differences in IVSA of MDPV in 96 h sessions (Nagy et al., 2023). No difference was observed between sexes in initial acquisition of MDPV IVSA in another study, however female rats obtained more infusions during a post-acquisition 5 week interval (Doyle et al., 2024).

The goal of this study was first to directly determine if there are sex-associated differences in the acquisition of IVSA of α-PVP or MA. Male and female groups were compared on mean infusions of drug obtained, as well as on the percentage of responses directed at the drug-associated lever, as dependent variables. A secondary goal was to analyze the proportion of animals that expressed a binge-like acquisition pattern identified previously for male rats self-administering MDPV and for female rats self-administering α-PVP (Aarde et al., 2015b; Javadi-Paydar et al., 2018a). The next primary goal was to determine if the per-infusion dose of MA could be adjusted to produce equivalent numbers of reinforcers obtained compared with the α-PVP groups. The impact of dose substitution under both Fixed Ratio and Progressive Ratio reward contingencies was next assessed for both drugs in all groups to determine the extent to which the training drug conferred any lasting differences in IVSA of the training dug or the alternate drug. This comparison maybe particularly important for fully assessing sex-differences, since a prior study showed no sex differences in oxycodone IVSA under a *PR* schedule in a study where males obtained fewer infusions in acquisition and a dose substitution under a *FR1* schedule of reinforcement (Nguyen et al., 2020). Finally, two α-PVP analogs that produce closely related pharmacological effects but differ in potency in IVSA, i.e., α-pyrrolidinopropiophenone HCl (α-PPP) and α-pyrrolidinohexiophenone HCl (α-PHP), were evaluated in FR and PR procedures to determine if the dose-effect relationships between these two compounds differed depending on sex or the initial training drug.

## 2. Materials and Methods

### 2.1 Subjects

Male (N=16) and Female (N=16) Wistar (CRL) were received at ∼12 weeks of age at the same time and housed in the same vivarium room throughout the study. The vivarium was kept on a 12:12 hour reversed light-dark cycle (lights out at 0800); studies were conducted during vivarium dark. Food and water were provided ad libitum in the home cage and in the experimental chambers. Procedures were conducted in accordance with protocols approved by the IACUC of the University of California, San Diego and were consistent with recommendations in the NIH Guide (Garber et al., 2011).

### 2.2 Drugs

The α-pyrrolidinopentiophenone HCl (α-PVP) and α-pyrrolidinohexiophenone HCl (α-PHP) were obtained from Cayman Chemical and d-methamphetamine HCl was provided by NIDA Drug Supply. The α-pyrrolidinopropiophenone HCl (α-PPP) was provided by Kenner C. Rice (NIDA Intramural Research Program). All drugs were dissolved in physiological saline for injection.

### 2.3 Surgery

Rats were surgically prepared with chronic indwelling intravenous catheters at 13-14 weeks of age using gas anesthesia and sterile procedures, as previously described (Aarde et al., 2015a; Nguyen et al., 2019; Nguyen et al., 2021). See **Supplementary Materials** for full description of implantation and catheter care. A minimum of 7 days was allowed for surgical recovery prior to starting the experiment. One female rat’s (α-PVP group) catheter port was chewed beyond repair by the cage mate prior to the start of the study. Exclusions due to catheter failure included female rats from the MA group after Sessions 10, 16, 25, 56, 57, female rats from the α-PVP group after Sessions 17, 75 and one male rat from the α-PVP group after Session 20. They were all retained as cage-mates for the animals remaining in the study. Animals with catheters that failed patency checks were discontinued from the study and any data that were collected after the previous passing of the test were excluded from analysis.

### 2.4 Intravenous Self-Administration (IVSA) Acquisition

At 18 weeks of age, IVSA was initiated in 2 h sessions with MA (0.05 mg/kg/infusion; N=8 per sex) or α-PVP (0.05 mg/kg/infusion; N=8 male, N=7 female) available on a Fixed Ratio 1 (FR 1) schedule of reinforcement, with sessions run on weekdays. If no response was emitted in the first 30 minutes of any IVSA session, a single non-contingent priming infusion was delivered. The female rats were run in one room and the male rats in a different room in two sequential daily cohorts per room. Drug identity groups were counterbalanced across the daily run cohorts. After 9 sessions, the per-infusion dose of MA was changed to 0.025 mg/kg for Sessions 10-21 to determine if doses leading to equivalent responding could be established. Consequently, the analysis of acquisition focused first on the initial 9 sessions and then on sessions 9-21 to assess the impact of the dose change.

### 2.5 IVSA Dose Substitution

#### 2.5.1 α-pyrrolidinopentiophenone (α-PVP) and methamphetamine (MA)

During Sessions 22-26 rats were tested with different per infusion doses of their training drug, α-PVP (0.0, 0.0125, 0.0250, 0.100, 0.300 mg/kg/infusion) or MA (0.0, 0.0125, 0.0250, 0.100, 0.300 mg/kg/infusion), in a counterbalanced order in 2 h sessions under the FR 1 schedule of reinforcement (starting on PND 159). Sessions 27-29 were baseline sessions, i.e., MA (0.025 mg/kg/infusion) or α-PVP (0.05 mg/kg/infusion) and then during Sessions 30-35 rats were tested in a similar dose-substitution procedure with the other drug. In Sessions 36-41, rats were maintained on 2 h sessions with the training drug switched, whereby the group originally trained on α-PVP received 0.025 mg/kg/infusion MA and the group originally trained on MA received 0.05 mg/kg/infusion α-PVP.

During Sessions 42-46, the groups originally trained on MA were tested with α-PVP (0.0, 0.0125, 0.0250, 0.100, 0.300 mg/kg/infusion) and groups originally trained on α-PVP assessed on MA (0.0, 0.0125, 0.0250, 0.100, 0.300 mg/kg/infusion) IVSA in a counterbalanced order in 3 h sessions under the Progressive Ratio schedule of reinforcement. In Sessions 47-51 this study was repeated except that the rats received doses of their original training drug in a counterbalanced order. In the PR paradigm, the required response ratio was increased after each reinforcer delivery, within a session as determined by the following equation (rounded to the nearest integer): Response Ratio=5e^(injection number^*^j)– 5 (Richardson and Roberts, 1996). In this study the j value was set to 0.2. Sessions were a maximum of 3 h in duration.

During Sessions 52-55 individual makeup sessions were conducted to replace sessions scheduled during the FR and PR dose substitutions which were disrupted by, e.g., failure of an infusion pump, inadvertent failure to start a session, tethers becoming disconnected, etc. This included four sessions for one male rat, three sessions for one female rat and one session for two male rats. The other rats were run in 3 hour sessions, FR1, with an 0.05 mg/kg/infusion dose of the original training drug during these sessions. Sessions 56-60 were continued additional baseline sessions with 0.05 mg/kg/infusion of each drug, for 3 h on Sessions 56-58 and 1 h on Sessions 59-60.

#### 2.5.2 α-pyrrolidinopropiophenone (α-PPP) and α-pyrrolidinohexiophenone (α-PHP)

The α-PPP analog has been shown to be less potent, and the α-PHP analog approximately equipotent, compared with α-PVP in pharmacological and self-administration assays in rats (Gannon et al., 2018a; Gannon et al., 2018b; Javadi-Paydar et al., 2018b). This comparison was included to further determine how group differences may, or may not, contribute to apparent behavioral potency and efficacy of drugs in IVSA. During Sessions 72-76 all remaining rats were tested with different per infusion doses of, α-PPP (0.0125, 0.0250, 0.050 0.100, 0.300 mg/kg/infusion), in a counterbalanced order in 1 h sessions under the FR 1 schedule of reinforcement. During Sessions 77-81 all remaining rats were tested with different per infusion doses of α-PHP (0.0125, 0.0250, 0.050, 0.100, 0.300 mg/kg/infusion), in a counterbalanced order in 1 h sessions under the FR 1 schedule of reinforcement. Session 82 was used to replace missing doses (N=2 male rats, N=3 female rats) from the α-PPP and α-PHP dose series. During Sessions 83-88 all remaining rats were tested with different per infusion doses of α-PPP and α-PHP (0.0125, 0.05, 0.300 mg/kg/infusion) available in a counterbalanced order under the PR schedule of reinforcement. Session 89 was used to replace 4 missing doses (N=4 male rats, N=5 female rats) from the α-PPP and α-PHP dose series.

### 2.6 Impact of Female Associated Odors on Male IVSA

Variability among prior reports on the presence or absence of sex differences in the IVSA of stimulant drugs is potentially influenced by a host of methodological choices. Dalla and colleagues warn of the *potential* practical logistical issue of conducting behavioral studies in both sexes (Dalla et al., 2024). The authors note that cleaning behavioral equipment is “*particularly important as the odors left behind on the apparatus from conspecifics may change the behavioral outcome*” (Dalla et al., 2024). This is coupled to a later caveat that tasks involving “*reward/motivation factors*” may “*overcome these effects*”. To determine if male rat IVSA was changed by the odor of female rats, an experiment was conducted in which female rats were switched to the male rat room and run in the boxes for one hour prior to the male session. No chamber cleaning or bedding changes were performed between the two sessions. This experiment was conducted over two separate days (Sessions 61 and 63) in ∼half of the male group each day because fewer female rats were left in the study at this stage, so it was necessary to use some female rats for more than one male evaluation. Sessions 64-71 were continued additional baseline sessions with 0.05 mg/kg/infusion of each drug in 1 h sessions.

### 2.7 Data Analysis

Infusions obtained, the percentage of responses directed at the drug-associated lever, total correct responses and breakpoints (for Progressive Ratio) were analyzed by ANOVA, or by mixed-effect models where there were missing values. Within-subjects factors of Session, and/or Dose were included where relevant. The between-subjects factor of Group was divided into Sex and Training Drug Identity factors in some analyses. In our approach a single priming infusion is delivered if no responses have been made within 30 minutes of session initiation under FR (this is not included for PR). These are rare after the initial few sessions of acquisition and are thus reported qualitatively where present, but not formally analyzed. The binge-like acquisition pattern was defined as obtaining 6 or more infusions in a single 5-minute bin during the session, based on prior studies (Aarde et al., 2015b; Javadi-Paydar et al., 2018a). One male rat from the MA group maintained catheter patency but exhibited 0-4 responses on sessions during the PR dose-substitutions involving all four drugs and was therefore excluded from all post-acquisition analyses. In all analyses, a criterion of P<0.05 was used to infer that a significant difference existed. Any significant main effects were followed with post-hoc analysis using Tukey (multi-level factors), Sidak (two-level factors) or Dunnett (to assess change relative to one of the treatment conditions.) correction. Initial three-factor analysis designed to test, e.g., the impact of group factors of Sex and Drug Identity, were followed with two-factor analysis (one Group factor) to facilitate the post-hoc analysis limited to orthogonal comparisons. All analysis used Prism for Windows (v. 10.3.0; GraphPad Software, Inc, San Diego CA).

## 3. Results

### 3.1 Acquisition

For the female groups, there were three priming infusions delivered in Sessions 1 and 2, two in Sessions 3-5, and 9, and one infusion in Sessions 6-8. One animal (MA) received a priming infusion on Sessions 2-9. Of these 17 primes, 11 were for MA animals (eight to the same individual) and 6 for α-PVP animals. For the male groups, there were four priming infusions delivered in Session 3, one priming infusion in Sessions 1, 2, 4, 6, and 9; no primes were delivered in Sessions 5, 7, and 8. Of these 9 primes, 7 were for MA animals (five to the same individual) and 2 for α-PVP animals.

The three-factor mixed-effects analysis confirmed there were significant effects of Session [F (8, 206) = 2.03; P<0.05] and of Drug identity [F (1, 26) = 16.11; P<0.0005] on reinforcers earned in the first 9 acquisition sessions (**Figure 1**). Therewere no significant effects of Sex or interactions with Sex confirmed. There were also significant effects of Session [F (8, 206) = 11.39; P<0.0001] and of Drug identity [F (1, 26) = 9.506; P<0.005] on the percentage of responses directed to the drug-associated lever. Altering the per-infusion dose of MA to 0.025 mg/kg/infusion resulted in an immediate increase in reinforcers obtained for the female and male groups. The female MA group approximated the mean of the α-PVP female group, however, the male MA group only increased mean infusions slightly and this only lasted for a few sessions. The mixed-effects analysis of sessions 9-21 confirmed there were significant effects of Session [F (12, 300) = 2.67; P<0.005], of Sex [F (1, 26) = 17.35; P<0.0005] and of Drug identity [F (1, 26) = 5.29; P<0.05] on reinforcers obtained. There were also significant effects of the interactions of Session with Sex [F (12, 300) = 2.27; P<0.05], and Sex with Drug identity [F (1, 26) = 5.83; P<0.05] confirmed. There were only significant effects of Sex [F (1, 26) = 5.19; P<0.05] and of Drug identity [F (1, 26) = 25.61; P<0.0001] on the percentage of responses directed at the drug-associated lever confirmed in the analysis.

**Figure 1:**
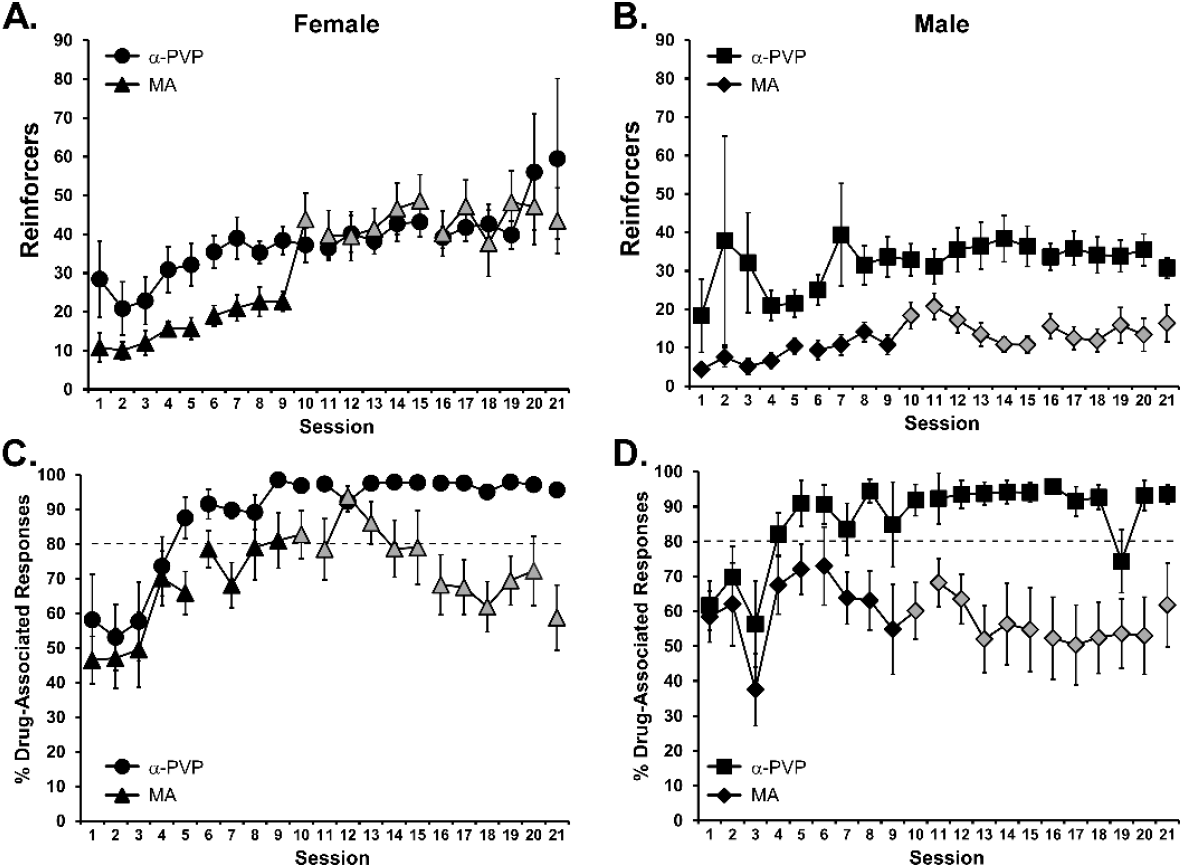
Mean (±SEM) drug reinforcers delivered (A, B) and percentage of responses directed to the drug-associated lever (C, D) obtained by groups of female rats (A, C) self-administering α-PVP (N=7; 0.05 mg/kg/infusion) or methamphetamine (N=7; 0.05 or 0.025 mg/kg/infusion) and by groups of male rats (B, D) self-administering α-PVP (N=8; 0.05 mg/kg/infusion) or methamphetamine (N=8; 0.05 or 0.025 mg/kg/infusion). Breaks in the series represent weekend breaks and gray symbols indicate where the methamphetamine dose was changed to 0.025 mg/kg/infusion.

### 3.2 Dose Substitution

#### 3.2.1 FR 1

##### 3.2.1.1 Training Drug

The number of infusions of the training drug obtained by rats in 2 h sessions under a Fixed Ratio 1 schedule of reinforcement depended on the available per-infusion dose (**Figure 2A**). The three-factor ANOVA confirmed there were significant effects of Sex [F (1, 22) = 20.33; P=0.0002], Drug [F (1, 22) = 44.57; P<0.0001], Dose [F (4, 88) = 84.03; P<0.0001] and the interaction of Dose with Drug [F (4, 88) = 24.05; P<0.0001] and Dose with Sex [F (4, 88) = 15.32; P<0.0001] on reinforcers acquired. There was no significant effect of the three-factor interaction [F (4, 88) = 2.35; P=0.0602].

**Figure 2:**
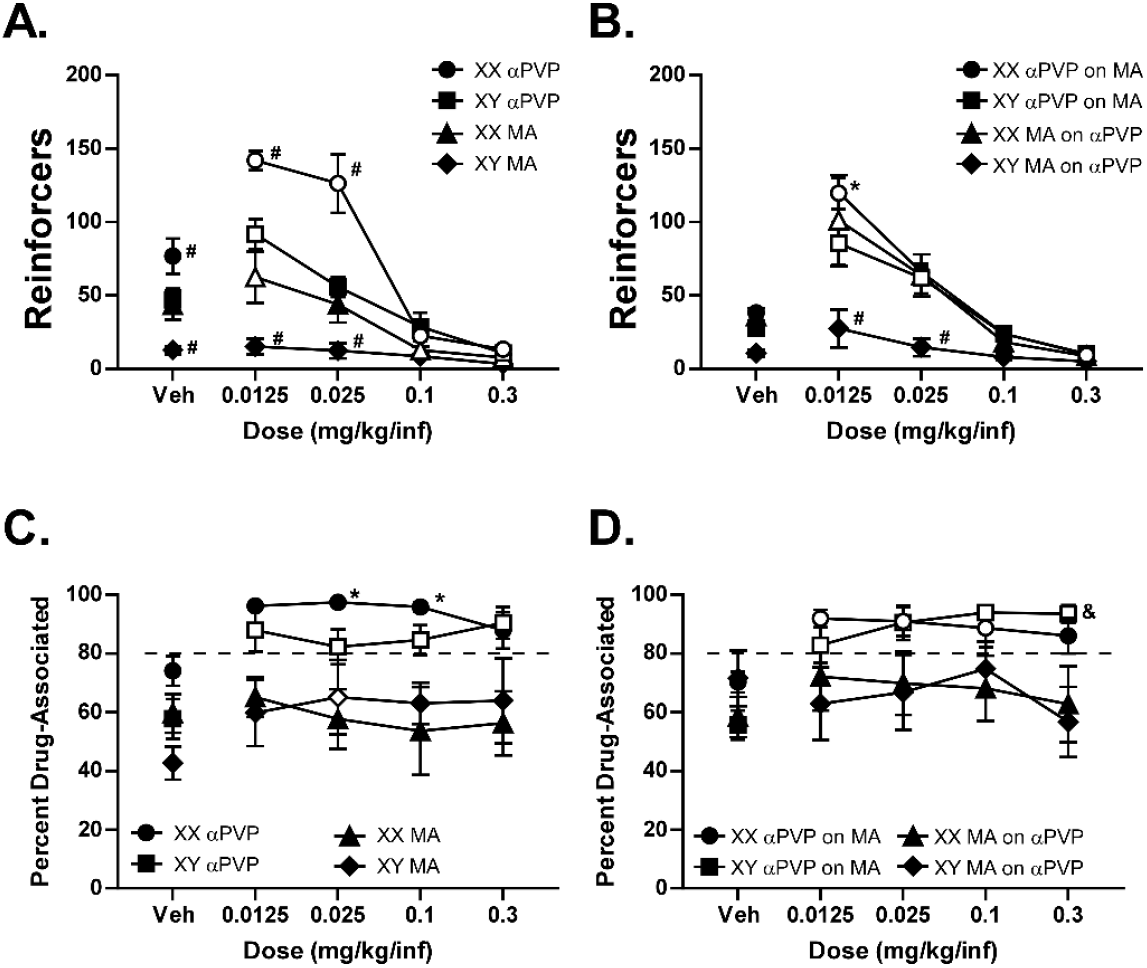
Mean (±SEM) infusions obtained (A, B) and the percent of responses directed to the drug-associated lever (C, D) by female (XX) α-PVP (N=6), male (XY) α-PVP (N=7), female MA (N=5-6; one animal failed to complete 0.1, 0.3 conditions) and male MA (N=8) groups responding under a Fixed Ratio 1 contingency in 2 hour sessions. The dotted line is included to emphasize when mean discrimination ratios exceed 80%. Data are depicted by original training drug (A, C) and when the alternate drug (B, D) was available. Bars indicate the SEM. Open symbols depict a significant difference from the Vehicle, within group. A significant difference from all other groups is indicated with #, a sex difference, within drug identity, with ^*^, and a difference between drugs, within sex, with &.

In the post-hoc test of orthogonal comparisons after the two-factor analysis, there were significant differences confirmed between all groups for the vehicle, 0.0125 and 0.025 doses, save that female MA and male α-PVP groups did not differ from each other at any of these doses. Within the female α-PVP group, there were significant differences of all active doses compared with vehicle, and between the 0.0125 and 0.025 doses and each of the 0.1 and 0.3 doses. Mean reinforcers for the male α-PVP group differed significantly between 0.0125 mg/kg/infusion and all other dose conditions including vehicle, between vehicle and 0.3 mg/kg/infusion and between the 0.025 dose and each of the 0.1 and 0.3 mg/kg/infusion doses. In the female MA group, the post-hoc confirmed significant differences between the 0.1 and 0.3 mg/kg/infusion doses compared with vehicle, 0.0125 and 0.025 mg/kg/infusion doses. There were no differences between doses confirmed for the male MA group.

The three-factor ANOVA confirmed there were significant effects of Drug [F (1, 22) = 14.75; P=0.001], and of Dose [F (4, 88) = 7.34; P<0.0001] on the <u>percentage of responses directed at the drug-associated lever</u> (**Figure 2C**). The post-hoc analysis of the marginal means confirmed that this was due to a lower percentage in the vehicle condition versus all active doses and a higher percentage in the female α-PVP group compared with both MA groups.

##### 3.2.1.2 Alternate Drug

When the alternate drug was available (**Figure 2B, D**), the number of reinforcers obtained again depended on dose. The three-factor ANOVA confirmed there were significant effects of Sex [F (1, 22) = 13.35; P=0.005], Drug [F (1, 22) = 8.86; P<0.01], Dose [F (4, 86) = 72.64; P<0.0001] and the interaction of Dose with Drug [F (4, 86) = 3.904; P<0.01], and Dose with Sex [F (4, 86) = 8.92; P<0.0001] on reinforcers acquired (**Figure 2B**).

When post-hoc test was limited to orthogonal comparisons after the two-factor analysis, there were significant differences confirmed by the post-hoc test between the male MA group (on α-PVP) and the other three groups for the 0.0125 and 0.025 doses. The male α-PVP group obtained fewer infusions of MA than did the female α-PVP group. Within the female α-PVP group, there were significant differences between 0.0125 and 0.3 mg/kg MA doses compared with vehicle, and between the 0.0125 and 0.025 doses and each of the 0.1 and 0.3 doses. Mean MA reinforcers obtained by the male α-PVP group in the 0.0125 and 0.025 mg/kg conditions differed significantly from each of the other conditions. Mean α-PVP reinforcers obtained by the female MA group in the 0.0125 and 0.025 mg/kg conditions differed significantly from each other and from each of the other conditions, save that vehicle did not differ from 0.025 mg/kg. There were no differences between doses confirmed for the male MA group self-administering α-PVP.

The three-factor ANOVA confirmed there were significant effects of Drug [F (1, 22) = 5.29; P=0.05], of Dose [F (4, 85) = 10.25; P<0.0001] as well as of the interaction of Dose with Drug [F (4, 85) = 7.88; P=0.0001] and of all three factors [F (4, 85) = 3.28; P=0.05] on the <u>percentage of responses directed at the drug-associated lever</u> (**Figure 2D**). The post-hoc analysis of the marginal means confirmed that this was due to a lower percentage in the vehicle condition versus all active doses but no differences between individual groups were confirmed. When post-hoc test was limited to orthogonal comparisons after the two-factor analysis, there was a significant difference between the male groups for the 0.3 mg/kg/infusion dose.

#### 3.2.2 PR

##### 3.2.2.1 Training Drug

The three-factor analysis of the IVSA of the original training drug confirmed significant effects of Dose [F (4, 82) = 12.54; P<0.0001], of Drug Identity [F (1, 21) = 4.96; P<0.05] and of the interaction of Dose with Drug Identity and Sex [F (4, 82) = 3.49; P<0.05] on the final completed ratio (**Figure 3A**). The post-hoc test confirmed that significantly higher final completed ratios were reached by male α-PVP (0.025, 0.1 mg/kg/infusion) and male MA (0.1 mg/kg/infusion) groups relative to the vehicle condition. The male α-PVP group achieved higher final completed ratios in the 0.1 mg/kg/infusion compared with the 0.0125 and 0.3 mg/kg/infusion conditions. The female α-PVP group reached higher final completed ratios in the 0.1 mg/kg/infusion condition compared with 0.0125 or 0.025 conditions. The female MA group reached significantly lower final completed ratios in the 0.3 mg/kg/infusion condition compared with 0.025 or 0.1 conditions. The only group difference confirmed was between the male and female MA groups in the 0.025 mg/kg/infusion condition.

**Figure 3:**
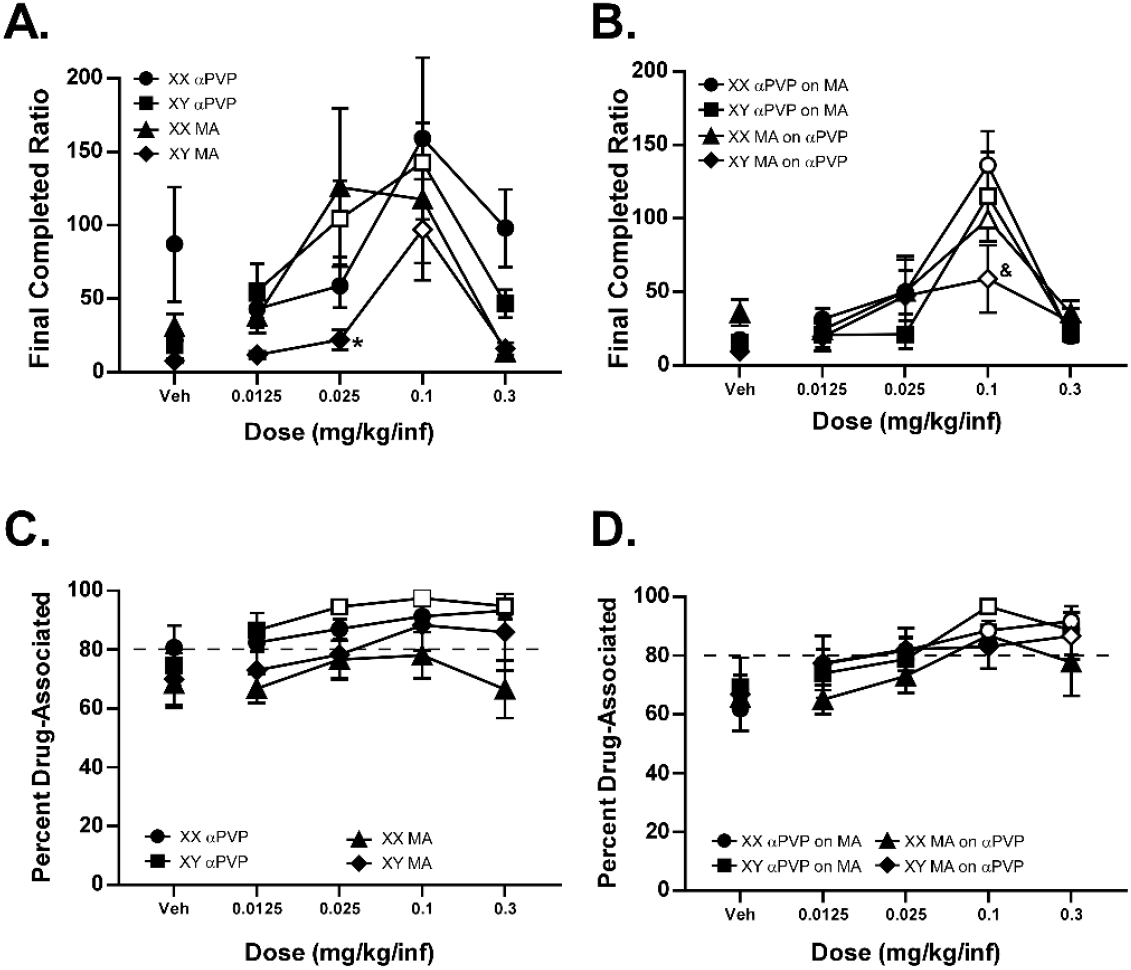
Mean (±SEM) final completed ratios (A, B) and the percent of responses directed to the drug-associated lever (C, D) by female (XX) α-PVP (N=6), male (XY) α-PVP (N=7), female MA (N=5-6; one animal failed to complete 0.1, 0.3 conditions) and male MA (N=7) groups. The dotted line is included to emphasize when mean discrimination ratios exceed 80%. Data are depicted by original training drug (A, C) and when the alternate drug (B, D) was available. Open symbols depict a significant difference from the <u>Vehicle</u>, within group. A significant sex difference, within drug identity, is indicated with ^*^ and a difference between drug groups, within sex, with &.

There were significant effects of Dose [F (4, 82) = 6.59; P<0.0005] and of Drug identity [F (1, 21) = 5.51; P<0.05] on the <u>percentage of responses directed at the drug-associated lever</u> (**Figure 3C**). The post-hoc test, collapsed across groups, confirmed that lever discrimination was lower in the vehicle condition compared with 0.025-0.3 mg/kg/infusion conditions, and lower in the 0.0125 mg/kg/infusion condition compared with the 0.1 mg/kg/infusion condition. The marginal mean analysis of the Drug identity factor confirmed significantly lower lever discrimination in the MA groups compared with the α-PVP groups.

##### 3.2.2.2 Alternate Drug

The three-factor analysis of the IVSA of the alternate drug confirmed significant effects of Dose [F (4, 83) = 26.81; P<0.0001] and of the interaction of Dose with Drug Identity [F (4, 83) = 3.495; P<0.05] on the final completed ratio (**Figure 3B**). The post-hoc test confirmed that significantly higher final completed ratios were reached by all groups in the 0.1 mg/kg/infusion condition relative to the vehicle condition. In addition, final completed ratios were significantly higher in the 0.1 mg/kg/infusion condition relative to all other conditions in each α-PVP-trained group and relative to the 0.0125 and 0.3 mg/kg/infusion conditions in the female MA-trained group.

There was only a significant effect of Dose [F (4, 83) = 13.50; P<0.0001] on the <u>percentage of responses directed at the drug-associated lever</u> (**Figure 3D**) confirmed in the analysis. The post-hoc test of the marginal means, collapsed across groups, confirmed that lever discrimination was lower in the vehicle condition compared with 0.025-0.3 mg/kg conditions, and lower in the 0.0125 mg/kg/infusion condition compared with the 0.1-0.3 mg/kg/infusion conditions.

#### 3.2.3 α-pyrrolidinopropiophenone (α-PPP) and α-pyrrolidinohexiophenone (α-PHP)

##### 3.2.3.1 Fixed Ratio

Responding depended on dose (0.0125, 0.0250, 0.050 0.100, 0.300 mg/kg/infusion) for α-PPP and α-PHP however the dose-effect curve differed between compounds (**Figure 4**). The highest numbers of infusions of α-PHP were obtained at the 0.0125 mg/kg/infusion dose, however the α-PPP analog engendered the most responding at the 0.05 mg/kg/infusion dose. A total of 8 female (N=3 from MA group) and 14 male (N=7 per training group) remained patent for these studies.

**Figure 4:**
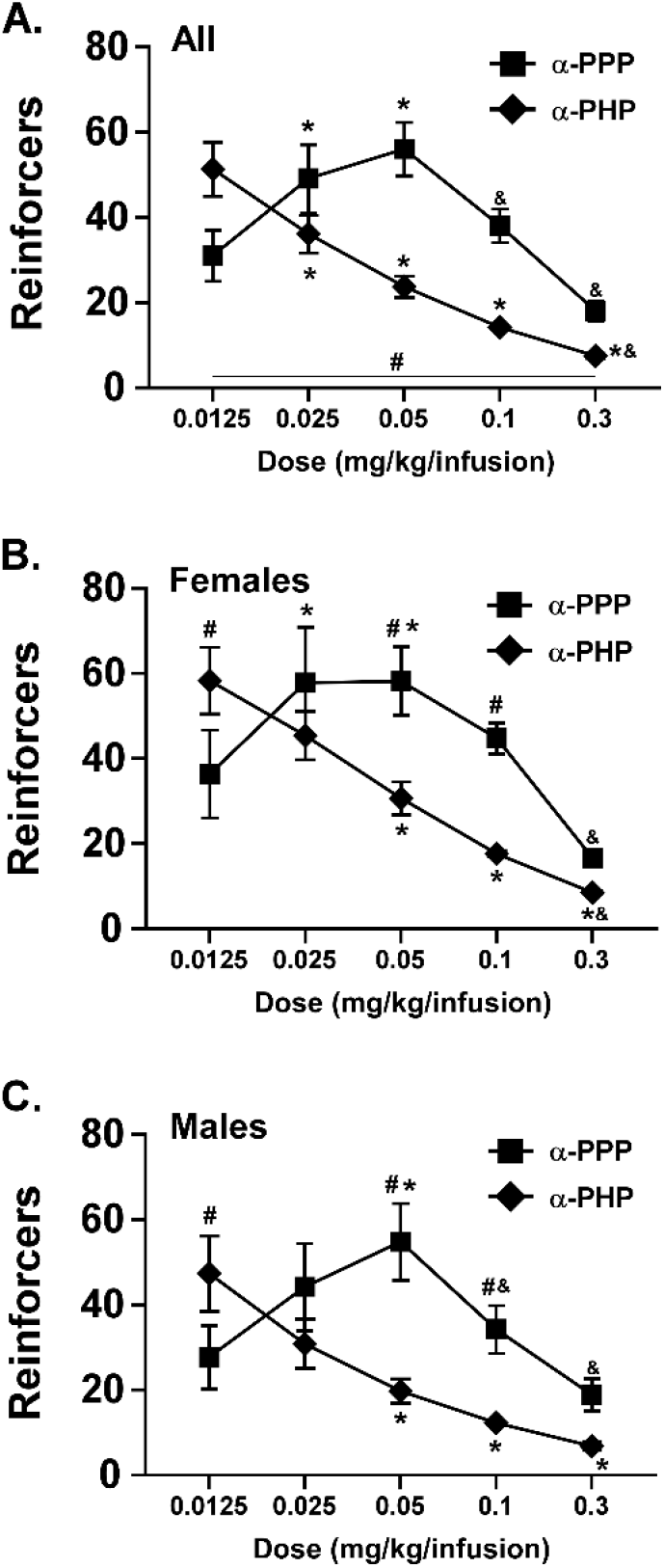
Mean (±SEM) reinforcers acquired for A) all, B) female (N=8) and C) male (N=14) rats completing the study of the IVSA of α-PPP and α-PHP under a FR schedule of reinforcement in 1 h sessions. A significant difference between α-PPP and α-PHP at a given dose is indicated with #, a difference from the 0.0125 mg/kg dose within drug with ^*^, and a difference from the 0.05 dose within drug with &.

The initial three-factor analysis combined original training drug groups within sex and confirmed significant effects of Dose [F (4, 80) = 24.47; P<0.0001], Drug Identity [F (1, 20) = 18.74; P<0.0005] and of the interaction of Dose with Drug Identity [F (4, 79) = 20.83; P<0.0001] on the number of reinforcers acquired; no significant effects of Sex were confirmed. The post-hoc analysis further confirmed that, collapsed across sex, significantly different numbers of reinforcers of α-PPP and α-PHP were obtained at each dose (**Figure 4A**). Furthermore, reinforcers acquired were significantly different from the 0.0125 mg/kg dose for α-PPP (0.025, 0.05 mg/kg) and α-PHP (0.025-0.3 mg/kg) and different from the 0.05 mg/kg dose for α-PPP (0.1, 0.3 mg/kg) and α-PHP (0.3 mg/kg). Within the female group a two-factor ANOVA confirmed significant effects of Dose [F (4, 56) = 17.58; P<0.0001], and of the interaction of Dose with Drug Identity [F (4, 56) = 7.11; P<0.0005] on the number of reinforcers acquired (**Figure 4B**). Within the male group a two-factor mixed-effects analysis confirmed significant effects of Dose [F (4, 103) = 13.68; P<0.0001], and of the interaction of Dose with Drug Identity [F (4, 103) = 10.60; P<0.0001] on the number of reinforcers acquired (**Figure 4C**). See **Supplemental Materials (Figure S6)** for analysis by training group.

##### 3.2.3.2 Progressive Ratio

The breakpoint (final completed ratio) reached in the PR study differed by dose and compound with the highest breakpoints achieved with the 0.3 mg/kg/infusion for α-PPP and the 0.05 mg/kg/infusion for α-PHP in each sex. One additional female (α-PVP-trained group) rat’s catheter lost patency during this study, thus N=7 for this analysis.

The three-factor analysis confirmed significant effects of Dose [F (2, 38) = 19.90; P<0.0001] and the interaction of Dose with Drug [F (2, 38) = 18.69; P<0.0001] on the final completed ratio; there were no significant effects of Sex (**Figure 5**). A follow-up two factor analysis collapsed across Sex confirmed significant effects of Dose [F (2, 40) = 22.99; P<0.0001], of Drug [F (1, 20) = 6.04; P<0.05] and the interaction of Dose with Drug [F (2, 40) = 19.03; P<0.0001] The post-hoc test confirmed that final completed ratios were higher for the 0.3 mg/kg/infusion α-PPP condition relative to the 0.0125 and 0.05 mg/kg/infusion conditions and higher for the 0.05 mg/kg/infusion PHP condition compared to the 0.0125 condition. Significantly higher ratios were also confirmed for α-PPP vs α-PHP for the 0.3 mg/kg/infusion conditions and higher for α-PHP vs α-PPP for the 0.05 mg/kg/infusion conditions. A follow-up analysis of the effect of training drug in the male groups did not confirm any significant contribution of training drug alone or in interaction with Dose or test Drug, see **Supplemental Materials, Figure S7**.

**Figure 5:**
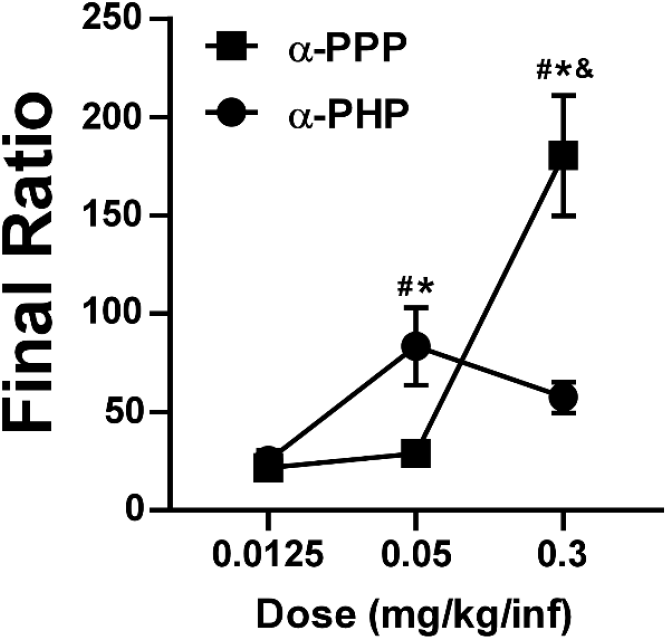
Mean (±SEM) final completed ratios for all female (N=7) and male (N=14) rats completing the study of the IVSA of α-PPP and α-PHP under a PR schedule of reinforcement. A significant difference between α-PPP and α-PHP is indicated with #, a difference from the 0.0125 mg/kg condition, within drug, with ^*^ and a difference from the 0.05 mg/kg condition, within drug, with &.

#### 3.2.4 Binge-like Acquisition Patterns

The Binge-like acquisition pattern was observed in 5/8 α-PVP males and 3/7 α-PVP females (see **Supplemental Materials, Figure S3**). Of the Binge acquisition α-PVP group, 7 completed the FR 1 and the PR dose-substitutions and of the No-Binge subgroup 6 completed each dose substitution. There was no significant effect of Binge / No-Binge grouping on reinforcers acquired [F (1, 11) = 1.69; P=0.2204] in the FR1 study, nor any significant effect of Binge grouping on the final completed ratio in the initial PR study [F (1, 11) = 0.47; P=0.5076] (see **Supplemental Materials, Figure S4**).

#### 3.2.5 Impact of female odor on male IVSA

There was no significant effect of session type (presence/absence of females in the operant chambers prior to running) on reinforcers acquired by male animals, although there was a significant effect of Drug identity confirmed (see **Supplemental Materials, Figure S5**).

## 4. Discussion

The study further confirms that α-pyrrolidinopentiophenone (α-PVP) is a highly effective reinforcer in rat intravenous self-administration (IVSA) paradigms, as has been previously demonstrated (Aarde et al., 2015a; Collins et al., 2019; Gannon et al., 2018a; Huskinson et al., 2017; Marusich et al., 2021; Xu et al., 2021). In this study both male and female α-PVP (0.05 mg/kg/infusion) groups attained stable intakes in 2 h sessions with drug-paired lever discrimination ratios above a group mean of 85% within 10 sessions; all individuals except one male exhibited ratios above 80% (**Figure 1**). In comparison, rats of each sex trained on methamphetamine (MA; 0.05 mg/kg/infusion) exhibited significantly lower levels of intake and lower drug-associated lever discrimination. In the case of the female animals, reducing the MA dose to 0.025 mg/kg/infusion on Session 10 produced an immediate increase in mean reinforcers obtained, to a level that matched the reinforcers obtained by the α-PVP rats. The male rats trained on MA obtained slightly more reinforcers in the following few sessions after the dose reduction, but quickly returned to about the same number of reinforcers obtained in the Sessions 8-9. Within each training drug, male and female rats self-administered the same number of reinforcers and reached similar discrimination ratios at the same rate and to the same degree with the original training dose of 0.05 mg/kg/infusion. This finding is at odds with some prior reports that female rats self-administer MA at higher rates (Roth and Carroll, 2004). However, sex differences *were* observed when the MA dose was decreased to attempt to match behavioral rates across drugs, within sex. This cautions that interpretation of sex differences, or lack thereof, with single training doses may fail to capture the complexity of how male and female rats self-administer a given drug. This may lead to an inappropriate general conclusion of either no sex difference, or that females self-administer more infusions, depending on the dose selected.

The training history affected self-administration in the dose substitution experiment under a Fixed-Ratio 1 (FR1) contingency. In particular, the male MA group obtained fewer infusions than all other groups at the lower two doses in both training-drug and alternate-drug substitutions (**Figure 2**). The male α-PVP group, conversely, self-administered MA at greater rates than the male MA group and in a pattern indistinguishable from their own training-drug substitution. The female groups’ self-administration was consistent with a combined effect of training history and drug identity, since the groups differed on their training drug but produced indistinguishable curves on the alternate drug (**Figure 2**). In the Progressive Ratio (**PR**) dose-substitutions the highest final completed ratio (breakpoint) was reached at the 0.1 mg/kg/infusion dose of each drug in all groups, save for female MA group where 0.025 and 0.1 mg/kg/infusion were similar. This further confirms a similarity of potency of the two drugs in dose-substitution procedures, even when there were apparent differences in the acquisition at the 0.05 mg/kg/infusion dose. The male MA-trained group reached lower final completed ratios than at least one other group for each drug, consistent with their FR1 dose-substitution. Our prior work has also shown that the reinforcing efficacy of a stimulant drug, as defined by infusions obtained in an IVSA experiment, depends on the drug with which the initial IVSA training has been conducted (Creehan et al., 2015; Vandewater et al., 2015). A full dose-substitution curve for MA IVSA in animals originally trained on 3,4-methylenedioxymethamphetamine (MDMA), pentylone or methylone shows that this is due to a difference in behavioral efficacy and not in potency, ruling out explanations based on pharmacodynamic tolerance from differential numbers of reinforcers acquired during acquisition (Khom et al., 2021).

We previously found that about half of individuals trained in the IVSA of α-PVP (female rats) or the closely related 3,4-methylenedioxypyrovalerone compound (male rats) exhibit a high-intake, binge-like session early in acquisition (Aarde et al., 2015b; Javadi-Paydar et al., 2018a). The ratio of binge/non-binge profiles observed in the combined male and female groups (8/15) is quite similar to our prior reports and there was no evidence of any major sex difference (**Supplemental Materials, Figure S3**). The results did not replicate the finding in male rats that binge phenotype animals self-administered more drug after acquisition (Aarde et al., 2015b) and confirmed a null impact of binge phenotype on the FR1 dose-substitution reported for female rats (Javadi-Paydar et al., 2018a).

The comparison of responding for the α-PPP and α-PHP analogs directly confirmed a potency difference that previously could only be inferred from indirect comparisons between studies. The α-PHP analog generated the highest rates of responding at the lowest dose (0.0125 mg/kg/infusion) in the FR procedure in 1 h sessions (**Figure 4**), similar to the dose-effect generated for α-PVP in 2 h sessions under a FR1 contingency (**Figure 2**). In contrast the highest rates of responding for α-PPP were at the 0.05 mg/kg/infusion dose in each sex. The highest rate of responding observed at the most efficacious dose of each compound was, however, quite similar. This suggests the α-PPP is of lower potency but of equal efficacy. The potency difference was further supported by the PR comparison whereby the animals reached higher breakpoints for the 0.3 mg/kg dose of α-PPP compared with the 0.3 mg/kg dose of α-PHP and vice versa at the 0.05 mg/kg/infusion dose. In this case the interpretation must be for the higher efficacy of α-PPP, although given the limited dose range evaluated this could also be due to a difference in the specific doses selected. (This latter interpretation relies on the assumption that some dose of α-PHP within the range that was not evaluated would be the most efficacious for PR.) There were no sex differences confirmed for either FR1 or PR comparisons of the α-PPP and α-PHP analogs. The male animals trained originally on α-PVP obtained more infusions across the dose ranges for each analog but the doses of α-PPP and α-PHP that generated the most responding were the same across MA and α-PVP trained groups (see **Supplemental Materials Figure S6**). Since only N=3 females trained on MA and N=4-5 females trained on α-PVP completed these studies, statistical assessment of sex differences based on original training group was not possible.

Female rats trained in the IVSA of cocaine who were then switched to MDPV IVSA did not develop into high-MDPV preferences that were observed in minor subset of animals trained originally with MDPV IVSA or food reinforcers (Doyle et al., 2021). Male Sprague-Dawley rats trained initially in MDPV IVSA obtained twice as many dug infusions compared with rats trained in cocaine IVSA using doses predicted to generate similar numbers of responses based on prior post-acquisition progressive ratio dose-substitution work (Seaman et al., 2022). When the drugs were switched, cocaine-trained animals obtained fewer infusions of MDPV than the MDPV-trained animals did during acquisition, but the MDPV-trained animals obtained similar amounts of cocaine compared with the cocaine-trained rats in acquisition. In the first FR1 dose-substitution in the present study, female rats trained on α-PVP obtained more infusions of MA than the MA-trained females did, and the MA-trained females obtained fewer infusions of α-PVP than the α-PVP trained group obtained, at the lower end of the dose-effect function.

The present study, and prior work (Seaman et al., 2022), emphasize that equating doses for IVSA across drugs based on the breakpoints obtained post-acquisition in a PR procedure may not lead to equivalent responding during initial acquisition under a FR1 schedule of reinforcement. Dose-effect functions generated by well-trained animals under FR1, such as the MA curve (**Figure 2B**) in the α-PVP-trained groups, likewise does not predict the impact of dose alterations during initial acquisition given the differential impact of changing to MA 0.025 mg/kg/infusion across male and female groups in this study.

The overall behavioral trajectory of the different groups in this study cautions against using operant response discrimination as conclusive evidence for/against the establishment of drug seeking behavior or as grounds for eliminating subjects from a study. Particularly under FR1 schedules, without any disincentive (e.g., an extended time-out for responses on the non-associated drug lever) there is very little real cost associated with low lever discrimination. It becomes an arbitrary and potentially misleading approach to exclude animals simply for low lever discrimination when they 1) continue to respond for drug infusions across scores of sessions, 2) substantially increase their drug-associated lever responding under increased work demands of a PR procedure and 3) exhibit dose sensitivity similar to other animals that do meet such arbitrary discrimination criteria. Finally, while 80% drug-associated lever responding is an individual exclusion criteria for some IVSA studies, others simply rely on showing statistical reliability of group mean responses on active versus inactive lever (see **Supplemental Materials Figure S2**) or a two-to-one ratio (Spencer et al., 2018; Stringfield et al., 2023). The field is better served by treating discrimination ratios as a dependent variable and not as a definition of self-administration in the sense of volitional drug-seeking behavior.

In summary this study showed that interpretations of sex-differences in stimulant self-administration depend on the training dose and that, more generally, antecedent training history is critical to assessing putative differences in potency or efficacy across stimulant drugs. It was also shown that within the α-pyrrolidino-phenone analogs, α-pyrrolidinopropiophenone is less potent than α-pyrrolidinopentiophenone or α-pyrrolidinohexiophenone, identifying a critical structural cutoff associated with length of the alkyl chain.

## Acknowledgements

These studies were supported by the United States Public Health Service NIH grant R01 DA042211 (MAT). The NIH/NIDA did not influence the study design, data interpretation, manuscript creation or the decision of when and what to publish from the studies conducted. The authors declare no competing financial interests.

## Literature Cited

Aarde, S.M., Creehan, K.M., Vandewater, S.A., Dickerson, T.J., Taffe, M.A., 2015a. In vivo potency and efficacy of the novel pyrrolidinopentiophenone cathinone alpha- and 3,4-methylenedioxypyrovalerone: self-administration and locomotor stimulation in male rats. Psychopharmacology 232(16), 3045–3055, doi: 10.1007/s00213-015-3944-8.

Aarde, S.M., Huang, P.K., Dickerson, T.J., Taffe, M.A., 2015b. Binge-like acquisition of 3,4-methylenedioxypyrovalerone (MDPV) self-administration and wheel activity in rats. Psychopharmacology 232(11), 1867–1877, doi: 10.1007/s00213-014-3819-4.

Albert, N., Catthoor, K., Morrens, M., 2022. Akathisia after chronic usage of synthetic cathinones: A case study. Front Psychiatry 13, 1046486, doi: 10.3389/fpsyt.2022.1046486.

Anker, J.J., Carroll, M.E., 2011. Females are more vulnerable to drug abuse than males: evidence from preclinical studies and the role of ovarian hormones. Current topics in behavioral neurosciences 8, 73–96, doi: 10.1007/7854_2010_93.

Barton, M., Wang, H., 2023. An Uncommon Presentation of Acute Thoracic Aortic Dissection. J Clin Med Res 15(6), 332–335, doi: 10.14740/jocmr4921.

Baumann, M.H., Partilla, J.S., Lehner, K.R., Thorndike, E.B., Hoffman, A.F., Holy, M., Rothman, R.B., Goldberg, S.R., Lupica, C.R., Sitte, H.H., Brandt, S.D., Tella, S.R., Cozzi, N.V., Schindler, C.W., 2013. Powerful cocaine-like actions of 3,4-Methylenedioxypyrovalerone (MDPV), a principal constituent of psychoactive ‘bath salts’ products. Neuropsychopharmacology : official publication of the American College of Neuropsychopharmacology 38(4), 552–562, doi: 10.1038/npp.2012.204.

Bender, C.M., Mao, C.E., Zangiabadi, A., 2022. Posterior Reversible Encephalopathy Syndrome With Hemorrhagic Conversion in a Patient With Active Polysubstance Abuse: A Case Report and Review of Literature. Cureus 14(10), e30909, doi: 10.7759/cureus.30909.

Brecht, M.L., Herbeck, D., 2014. Time to relapse following treatment for methamphetamine use: a long-term perspective on patterns and predictors. Drug and alcohol dependence 139, 18–25, doi: 10.1016/j.drugalcdep.2014.02.702.

CBS17, 2020. Raleigh man charged with possession of drug known as ‘monkey dust’ CBS17.COM. CBS17, Raleigh, NC.

Clayton, J.A., Collins, F.S., 2014. Policy: NIH to balance sex in cell and animal studies. Nature 509(7500), 282–283, doi:

Collins, G.T., Sulima, A., Rice, K.C., France, C.P., 2019. Self-administration of the synthetic cathinones 3,4-methylenedioxypyrovalerone (MDPV) and alpha-pyrrolidinopentiophenone (alpha-PVP) in rhesus monkeys. Psychopharmacology 236(12), 3677–3685, doi: 10.1007/s00213-019-05339-4.

Creehan, K.M., Vandewater, S.A., Taffe, M.A., 2015. Intravenous self-administration of mephedrone, methylone and MDMA in female rats. Neuropharmacology 92, 90–97, doi: 10.1016/j.neuropharm.2015.01.003.

Dalla, C., Jaric, I., Pavlidi, P., Hodes, G.E., Kokras, N., Bespalov, A., Kas, M.J., Steckler, T., Kabbaj, M., Wurbel, H., Marrocco, J., Tollkuhn, J., Shansky, R., Bangasser, D., Becker, J.B., McCarthy, M., Ferland-Beckham, C., 2024. Practical solutions for including sex as a biological variable (SABV) in preclinical neuropsychopharmacological research. Journal of neuroscience methods 401, 110003, doi: 10.1016/j.jneumeth.2023.110003.

DEA, 2014. National Forensic Laboratory Information System: Midyear Report 2013. https://www.nflis.deadiversion.usdoj.gov/.

DEA, 2019. National Forensic Laboratory Information System: Drug 2018 Annual Report. https://www.nflis.deadiversion.usdoj.gov/.

DEA, 2020. National Forensic Laboratory Information System: Drug 2019 Annual Report. https://www.nflis.deadiversion.usdoj.gov/.

DEA, 2021. National Forensic Laboratory Information System: snapshot December 2020. https://www.nflis.deadiversion.usdoj.gov/.

DEA, 2024. National Forensic Laboratory Information System: snapshot December 2024. https://www.nflis.deadiversion.usdoj.gov/publicationsRedesign.xhtml.

Doyle, M.R., Beltran, N.M., Bushnell, M.S.A., Syed, M., Acosta, V., Desai, M., Rice, K.C., Serafine, K.M., Gould, G.G., Daws, L.C., Collins, G.T., 2024. Effects of access condition on substance use disorder-like phenotypes in male and female rats self-administering MDPV or cocaine. bioRxiv, doi: 10.1101/2024.03.04.583431.

Doyle, M.R., Sulima, A., Rice, K.C., Collins, G.T., 2021. MDPV self-administration in female rats: influence of reinforcement history. Psychopharmacology 238(3), 735–744, doi: 10.1007/s00213-020-05726-2.

Enabi, J., Shah, K., Kondakindi, H., Mukkera, S., 2024. A Rare Presentation of Polyarteritis Nodosa (PAN). Cureus 16(2), e55143, doi: 10.7759/cureus.55143.

Fox12Staff, 2019. DOJ: Vancouver man arrested for trafficking ‘bath salts’ out of mobile home, storage units Fox 12 Oregon. KPTV-KPDX Broadcasting Corporation, Beaverton, OR.

Funke, J.R., Hwang, E.K., Wunsch, A.M., Baker, R., Engeln, K.A., Murray, C.H., Milovanovic, M., Caccamise, A.J., Wolf, M.E., 2023. Persistent Neuroadaptations in the Nucleus Accumbens Core Accompany Incubation of Methamphetamine Craving in Male and Female Rats. eNeuro 10(3), doi: 10.1523/ENEURO.0480-22.2023.

Gannon, B.M., Baumann, M.H., Walther, D., Jimenez-Morigosa, C., Sulima, A., Rice, K.C., Collins, G.T., 2018a. The abuse-related effects of pyrrolidine-containing cathinones are related to their potency and selectivity to inhibit the dopamine transporter. Neuropsychopharmacology : official publication of the American College of Neuropsychopharmacology 43(12), 2399–2407, doi: 10.1038/s41386-018-0209-3.

Gannon, B.M., Galindo, K.I., Mesmin, M.P., Sulima, A., Rice, K.C., Collins, G.T., 2018b. Relative reinforcing effects of second-generation synthetic cathinones: Acquisition of self-administration and fixed ratio dose-response curves in rats. Neuropharmacology 134(Pt A), 28–35, doi: 10.1016/j.neuropharm.2017.08.018.

Gannon, B.M., Rice, K.C., Collins, G.T., 2017. Reinforcing effects of abused ‘bath salts’ constituents 3,4-methylenedioxypyrovalerone and alpha-pyrrolidinopentiophenone and their enantiomers. Behavioural pharmacology 28(7), 578–581, doi: 10.1097/FBP.0000000000000315.

Garber, J.C., Barbee, R.W., Bielitzki, J.T., Clayton, L.A., Donovan, J.C., Hendriksen, C.F.M., Kohn, D.F., Lipman, N.S., Locke, P.A., Melcher, J., Quimby, F.W., Turner, P.V., Wood, G.A., Wurbel, H., 2011. Guide for the Care and Use of Laboratory Animals, 8th Edition. National Academies Press, Washington D.C.

Han, B., Cotto, J., Etz, K., Einstein, E.B., Compton, W.M., Volkow, N.D., 2021. Methamphetamine Overdose Deaths in the US by Sex and Race and Ethnicity. JAMA Psychiatry, doi: 10.1001/jamapsychiatry.2020.4321.

Holsman, M., 2021. St. Lucie County sheriff’s probe results in seizure of guns, arrests, TC Palm. Treasure Coast Newspapers, Fort Pierce, FL.

Huskinson, S.L., Naylor, J.E., Townsend, E.A., Rowlett, J.K., Blough, B.E., Freeman, K.B., 2017. Self-administration and behavioral economics of second-generation synthetic cathinones in male rats. Psychopharmacology 234(4), 589–598, doi: 10.1007/s00213-016-4492-6.

Javadi-Paydar, M., Harvey, E.L., Grant, Y., Vandewater, S.A., Creehan, K.M., Nguyen, J.D., Dickerson, T.J., Taffe, M.A., 2018a. Binge-like acquisition of alpha-pyrrolidinopentiophenone (alpha-PVP) self-administration in female rats. Psychopharmacology 235(8), 2447–2457, doi: 10.1007/s00213-018-4943-3.

Javadi-Paydar, M., Nguyen, J.D., Vandewater, S.A., Dickerson, T.J., Taffe, M.A., 2018b. Locomotor and reinforcing effects of pentedrone, pentylone and methylone in rats. Neuropharmacology 134(Pt A), 57–64, doi: 10.1016/j.neuropharm.2017.09.002.

Khom, S., Nguyen, J.D., Vandewater, S.A., Grant, Y., Roberto, M., Taffe, M.A., 2021. Self-Administration of Entactogen Psychostimulants Dysregulates Gamma-Aminobutyric Acid (GABA) and Kappa Opioid Receptor Signaling in the Central Nucleus of the Amygdala of Female Wistar Rats. Front Behav Neurosci 15, 780500, doi: 10.3389/fnbeh.2021.780500.

Kolanos, R., Sakloth, F., Jain, A.D., Partilla, J.S., Baumann, M.H., Glennon, R.A., 2015. Structural Modification of the Designer Stimulant alpha-Pyrrolidinovalerophenone (alpha-PVP) Influences Potency at Dopamine Transporters. ACS chemical neuroscience 6(10), 1726–1731, doi: 10.1021/acschemneuro.5b00160.

Lin, C.H., Chen, J.J., Chan, C.H., 2023. Comparison of Psychiatric and Clinical Profiles Between People Who Use Synthetic Cathinones and Methamphetamine: A Matched Case-Control Study. Journal of clinical psychopharmacology 43(2), 122-130, doi: 10.1097/JCP.0000000000001649.

Marusich, J.A., Gay, E.A., Watson, S.L., Blough, B.E., 2021. Alpha-pyrrolidinopentiophenone and mephedrone self-administration produce differential neurochemical changes following short-or long-access conditions in rats. Eur J Pharmacol 897, 173935, doi: 10.1016/j.ejphar.2021.173935.

Miller, A.E., Daiwile, A.P., Cadet, J.L., 2022. Sex-Dependent Alterations in the mRNA Expression of Enzymes Involved in Dopamine Synthesis and Breakdown After Methamphetamine Self-Administration. Neurotoxicity research 40(5), 1464–1478, doi: 10.1007/s12640-022-00545-z.

Mohr, A.L.A., Friscia, M., Yeakel, J.K., Logan, B.K., 2018. Use of synthetic stimulants and hallucinogens in a cohort of electronic dance music festival attendees. Forensic science international 282, 168–178, doi: 10.1016/j.forsciint.2017.11.017.

Nagy, E.K., Leyrer-Jackson, J.M., Hood, L.E., Acuna, A.M., Olive, M.F., 2023. Effects of repeated binge intake of the pyrovalerone cathinone derivative 3,4-methylenedioxypyrovalerone on prefrontal cytokine levels in rats - a preliminary study. Front Behav Neurosci 17, 1275968, doi: 10.3389/fnbeh.2023.1275968.

Nash, J., 2024. Multiple arrests made by agencies working together, Lake Okeechobee News. LakeONews.com, Okeechobee, FL.

Nguyen, J.D., Creehan, K.M., Kerr, T.M., Taffe, M.A., 2020. Lasting effects of repeated Δ(9) - tetrahydrocannabinol vapour inhalation during adolescence in male and female rats. British journal of pharmacology 177(1), 188–203, doi: 10.1111/bph.14856.

Nguyen, J.D., Grant, Y., Creehan, K.M., Hwang, C.S., Vandewater, S.A., Janda, K.D., Cole, M., Taffe, M.A., 2019. Delta(9)-tetrahydrocannabinol attenuates oxycodone self-administration under extended access conditions. Neuropharmacology 151, 127–135, doi: 10.1016/j.neuropharm.2019.04.010.

Nguyen, J.D., Grant, Y., Taffe, M.A., 2021. Paradoxical changes in brain reward status during oxycodone self-administration in a novel test of the negative reinforcement hypothesis. British journal of pharmacology, doi: 10.1111/bph.15520.

Reichel, C.M., Chan, C.H., Ghee, S.M., See, R.E., 2012. Sex differences in escalation of methamphetamine self-administration: cognitive and motivational consequences in rats. Psychopharmacology 223(4), 371–380, doi: 10.1007/s00213-012-2727-8.

Richardson, N.R., Roberts, D.C.S., 1996. Progressive ratio schedules in drug self-administration studies in rats: a method to evaluate reinforcing efficacy. Journal of neuroscience methods 66, 1–11, doi:

Roth, M.E., Carroll, M.E., 2004. Sex differences in the acquisition of IV methamphetamine self-administration and subsequent maintenance under a progressive ratio schedule in rats. Psychopharmacology 172(4), 443–449, doi:

SAMHSA, 2023. Key Substance Use and Mental Health Indicators in the United States: Results from the 2022 National Survey on Drug Use and Health, in: CBHSQ (Ed.). Substance Abuse and Mental Health Services Administration, Rockville, MD.

Seaman, R.W., Jr., Rice, K.C., Collins, G.T., 2022. Relative reinforcing effects of cocaine and 3,4-methylenedioxypyrovalerone (MDPV) under a concurrent access self-administration procedure in rats. Drug and alcohol dependence 232, 109299, doi: 10.1016/j.drugalcdep.2022.109299.

Shansky, R.M., Murphy, A.Z., 2021. Considering sex as a biological variable will require a global shift in science culture. Nat Neurosci, doi: 10.1038/s41593-021-00806-8.

Simmler, L., Buser, T., Donzelli, M., Schramm, Y., Dieu, L.H., Huwyler, J., Chaboz, S., Hoener, M., Liechti, M., 2013. Pharmacological characterization of designer cathinones in vitro. British journal of pharmacology 168(2), 458–470, doi: 10.1111/j.1476-5381.2012.02145.x.

Spencer, S., Neuhofer, D., Chioma, V.C., Garcia-Keller, C., Schwartz, D.J., Allen, N., Scofield, M.D., Ortiz-Ithier, T., Kalivas, P.W., 2018. A Model of Delta(9)-Tetrahydrocannabinol Self-administration and Reinstatement That Alters Synaptic Plasticity in Nucleus Accumbens. Biol Psychiatry 84(8), 601–610, doi: 10.1016/j.biopsych.2018.04.016.

Stringfield, S.J., Sanders, B.E., Suppo, J.A., Sved, A.F., Torregrossa, M.M., 2023. Nicotine Enhances Intravenous Self-administration of Cannabinoids in Adult Rats. Nicotine Tob Res 25(5), 1022–1029, doi: 10.1093/ntr/ntac267.

Vandewater, S.A., Creehan, K.M., Taffe, M.A., 2015. Intravenous self-administration of entactogen-class stimulants in male rats. Neuropharmacology 99, 538–545, doi: 10.1016/j.neuropharm.2015.08.030.

Webb, P., Ireland, J., Colledge-Frisby, S., Peacock, A., Leung, J., Vickerman, P., Farrell, M., Hickman, M., Grebely, J., Degenhardt, L., 2024. Patterns of drug use among people who inject drugs: A global systematic review and meta-analysis. The International journal on drug policy 128, 104455, doi: 10.1016/j.drugpo.2024.104455.

Xu, P., Lai, M., Fu, D., Liu, H., Wang, Y., Shen, H., Zhou, W., 2021. Reinforcing and discriminative-stimulus effects of two pyrrolidine-containing synthetic cathinone derivatives in rats. Pharmacology, biochemistry, and behavior 203, 173128, doi: 10.1016/j.pbb.2021.173128.

Zlebnik, N.E., Holtz, N.A., Lepak, V.C., Saykao, A.T., Zhang, Y., Carroll, M.E., 2021. Age-specific treatment effects of orexin/hypocretin-receptor antagonism on methamphetamine-seeking behavior. Drug and alcohol dependence 224, 108719, doi: 10.1016/j.drugalcdep.2021.108719.

