## Supplemental Materials for "Methamphetamine and α-pyrrolidinopentiophenone (α-PVP) Intravenous Self-Administration in Female and Male Rats"

### 2. Supplementary Materials and Methods

Surgery:

Rats were surgically prepared with chronic indwelling intravenous catheters at 13-14 weeks of age using gas anesthesia and sterile procedures. Catheters consisted of a 14-cm length polyurethane based tubing (MicroRenathane®, Braintree Scientific, Inc, Braintree MA, USA) fitted to a guide cannula (Plastics one, Roanoke, VA) curved at an angle and encased in dental cement anchored to an ~3-cm circle of durable mesh. The catheter tubing was passed subcutaneously from a port at the back, inserted into the jugular vein and secured gently with suture thread. A liquid tissue adhesive was used to close the incisions (3M™ Vetbond™ Tissue Adhesive; 1469S B). Catheters were flushed with ~0.2-0.3 ml heparinized (166.7 USP/ml) saline before sessions and ~0.2-0.3 ml heparinized saline containing cefazolin (100 mg/mL) after sessions. Catheter patency was assessed with the administration of ~0.2 ml (10 mg/ml) of the ultra-short-acting barbiturate anesthetic, Brevital sodium (1 % methohexital sodium; Eli Lilly, Indianapolis, IN), i.v.. Animals with patent catheters exhibit pronounced loss of muscle tone within ~3 s of infusion. Animals that failed to display these signs were discontinued from the study and any data that were collected after the previous passing of the test were excluded from analysis.

Data Analysis:

Body weight, infusions obtained, the percentage of responses directed at the drug-associated lever, the number of responses directed at the drug-associated and alternate levers, total correct responses and breakpoints (for Progressive Ratio) were analyzed by ANOVA, or by mixed-effect models where there were missing values. Within-subjects factors of Group, Session, Dose and/or Lever were included where relevant; Group was divided into Sex and Training Drug Identity in some analyses. The binge-like acquisition pattern was defined as obtaining 6 or more infusions in a single 5 minute bin during the session, based on prior studies (Aarde et al., 2015; Javadi-Paydar et al., 2018). One male rat from the MA group maintained catheter patency but exhibited 0-4 responses on sessions during the PR dose-substitutions involving all four drugs and was therefore excluded from all post-acquisition analyses.

In all analyses, a criterion of $P<0.05$ was used to infer that a significant difference existed. Any significant main effects were followed with post-hoc analysis using Tukey (multi-level factors), Sidak (two-level factors) or Dunnett (to assess change relative to one of the Treatment Conditions.) correction. Initial three-factor analysis designed to test, e.g., the impact of group factors of Sex and Drug Identity, were followed with two-factor analysis (one Group factor) to facilitate the post-hoc analysis limited to orthogonal comparisons. All analysis used Prism for Windows (v. 10.3.0; GraphPad Software, Inc, San Diego CA).

### 3. Supplementary Results

### Bodyweight:

There was no effect of initial training drug on the body weight (**Figure S1**) of the female [Group: $F(1, 12) = 0.13$; $P=0.73$, Week: $F(3.341, 40.10) = 65.55$; $P<0.0001$, Interaction: $F(27, 324) = 0.32$; $P=0.9996$] or the male [Group: $F(1, 14) = 0.05$; $P=0.82$, Week: $F(1.95, 26.59) = 218.9$; $P<0.0001$, Interaction: $F(27, 368) = 0.50$; $P=0.98$] rats. Two female rats were excluded from this analysis as they were not in the self-administration study after day 10. A male rat was found dead after week 31.

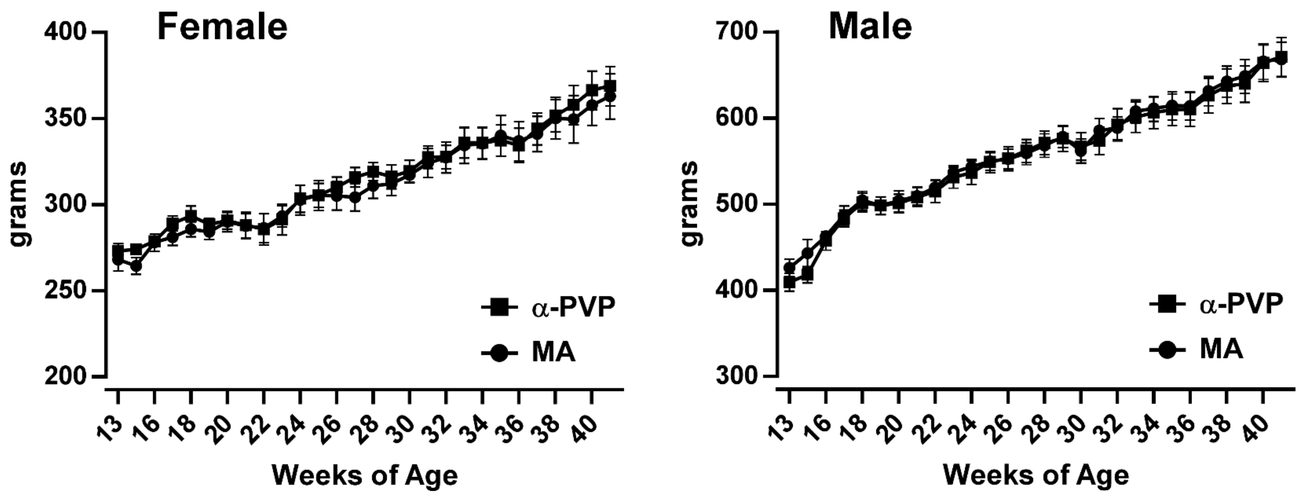


***Figure S1:*** *Mean (±SEM) body weight of the female and male rats trained on alpha-PVP and methamphetamine.*

### Drug-Associated and Alternate Lever Reponses During Acquisition:

Although the ratio of responses on the drug-associated lever is often used as a criterion for acquisition of self-administration behavior, there are reports which prefer to use an inferential statistical comparison of responses on the drug-associated and the non-associated lever. This analysis was conducted over the first 21 days for each of the four groups using all responses on the Drug-Associated and the alternate lever (**Figure S2**), including during the Time Out interval. Individual responses for several sessions are excluded from these graphs due to unusually higher numbers of responses on the drug-associated lever, including one α-PVP female (Sessions 20, 20; 1290-5193), one α-PVP male (Sessions 2, 3 and 7; 810-2311). Equipment error is not likely the cause for the former, since another female animal run in the same operant chamber did not exhibit this unusual high response pattern on those two days. The session for the male rat (#328) was associated with the binge-like intake pattern, see below. There were significantly more responses on the drug-associated lever for α-PVP female [Session: n.s., Lever: F (1.000, 6.000) = 15.44; P<0.01, Interaction: n.s.], MA female [Session: F (2.765, 16.59) = 5.301; P<0.05, Lever: F (1.000, 6.000) = 25.96; P<0.005, Interaction: n.s.], α-PVP male [Session: n.s.; Lever: F (1.000, 7.000) = 35.05; P<0.001, Interaction: n.s.] rats, but not for the MA male rats [Session: F (3.067, 21.47) = 3.761; P=0.0253 Lever: n.s., Interaction: n.s.].

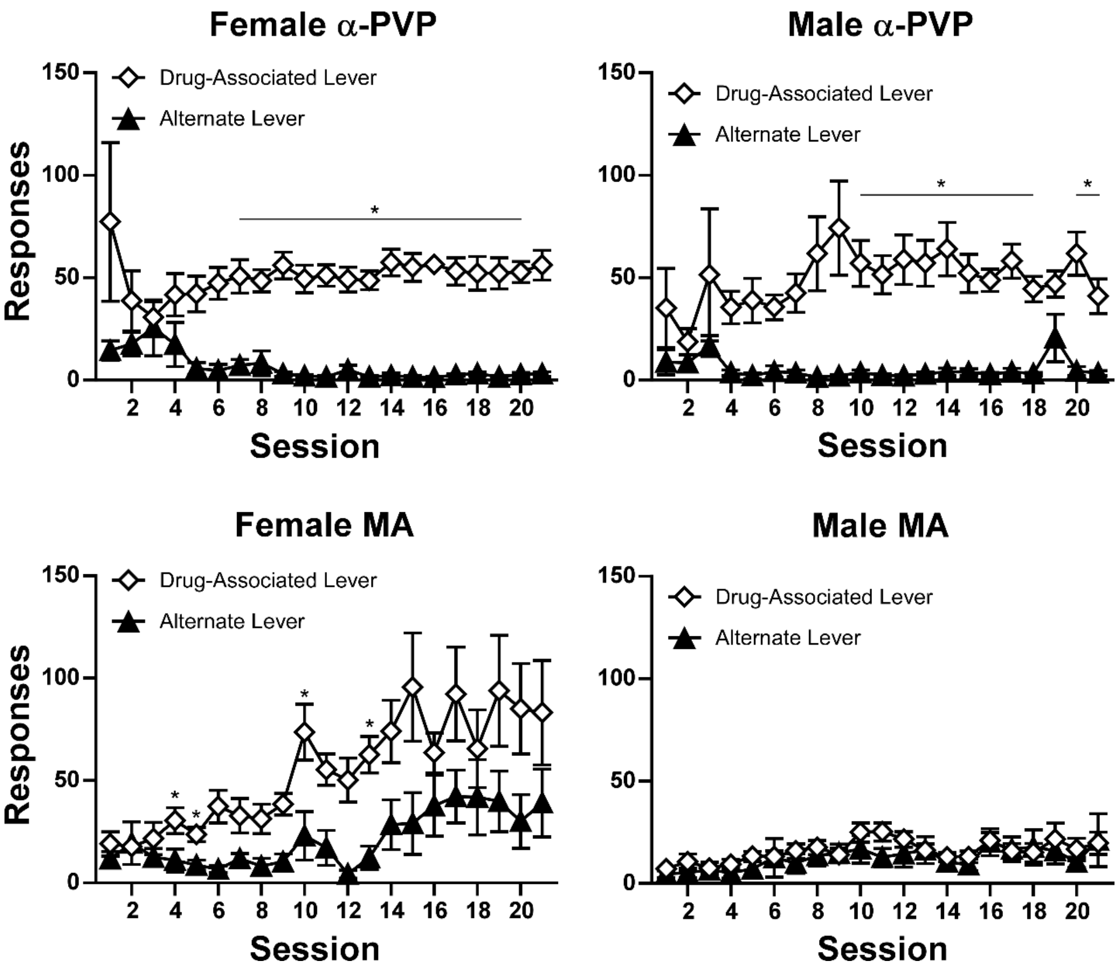


***Figure S2:*** *Mean (±SEM) responses on the drug-associated and alternate levers for the female and male rats trained on alpha-PVP and methamphetamine. A significant difference between levers is indicated with *.*

### Binge-like Acquisition Patterns:

The Binge-like acquisition pattern was observed in 5 of 8 α-PVP males and 3 of 7 α-PVP females (**Figure S3**). Although the pattern for female rat #332 was similar to the binge phenotype, this animal generated only several 5 infusion bins on day 4, but no 6 infusion bins, and thus did not meet the *a priori* criterion.

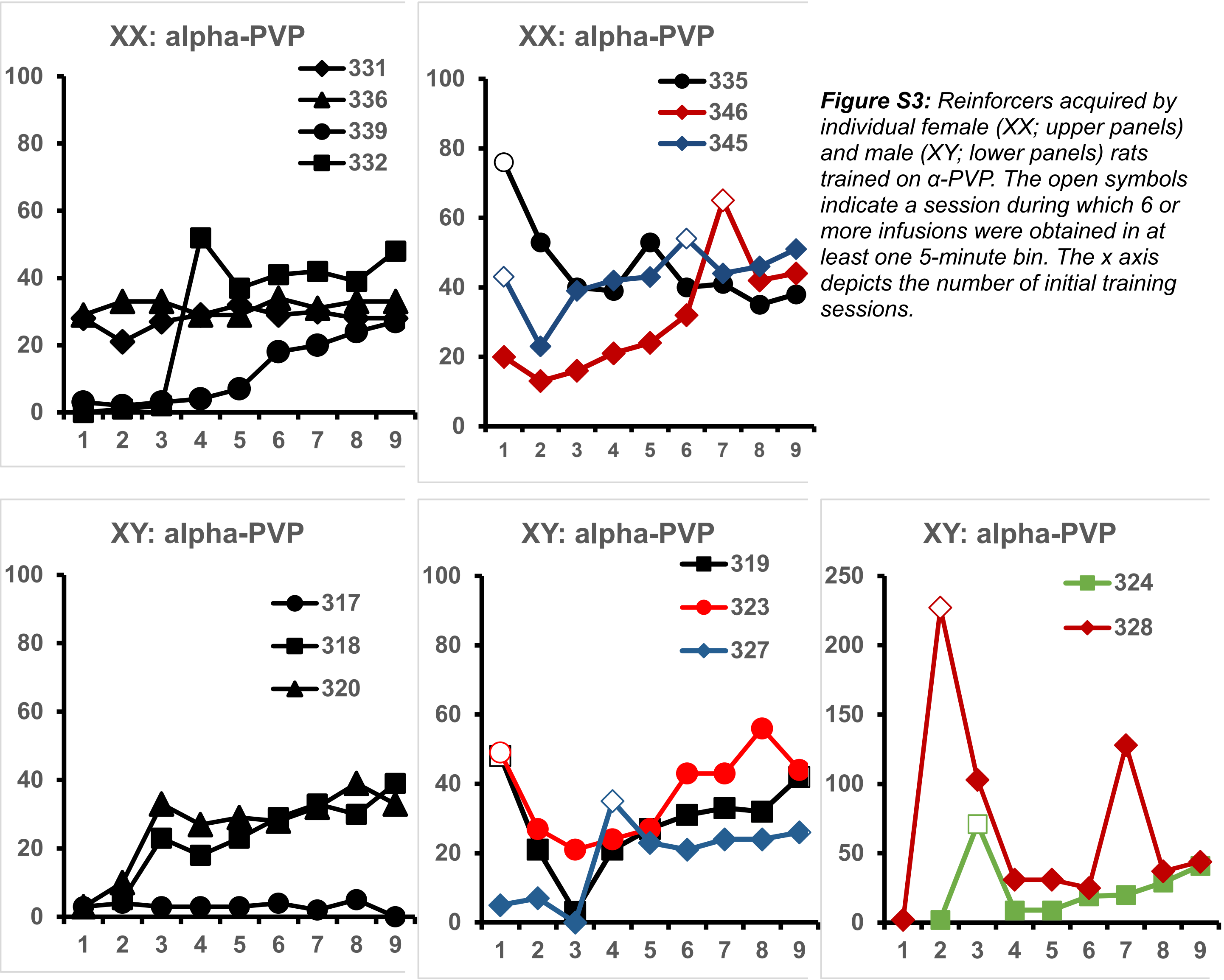


***Figure S3:*** *Reinforcers acquired by individual female (XX; upper panels) and male (XY; lower panels) rats trained on α-PVP. The open symbols indicate a session during which 6 or more infusions were obtained in at least one 5-minute bin. The x axis depicts the number of initial training sessions.*

### Dose Substitution by Acquisition Binge-like Phenotype:

Of the Binge acquisition α-PVP rats, 7 completed the FR 1 and the PR dose-substitutions and of the No-Binge subgroup 6 completed each dose substitution (**Figure S4**). There was no significant effect of Binge / Non-Binge grouping on reinforcers acquired [$F(1, 11) = 1.69$; $P=0.2204$] in the FR1 study, nor any significant effect of Binge grouping on the final completed ratio in the PR study [$F(1, 11) = 0.47$; $P=0.5076$].

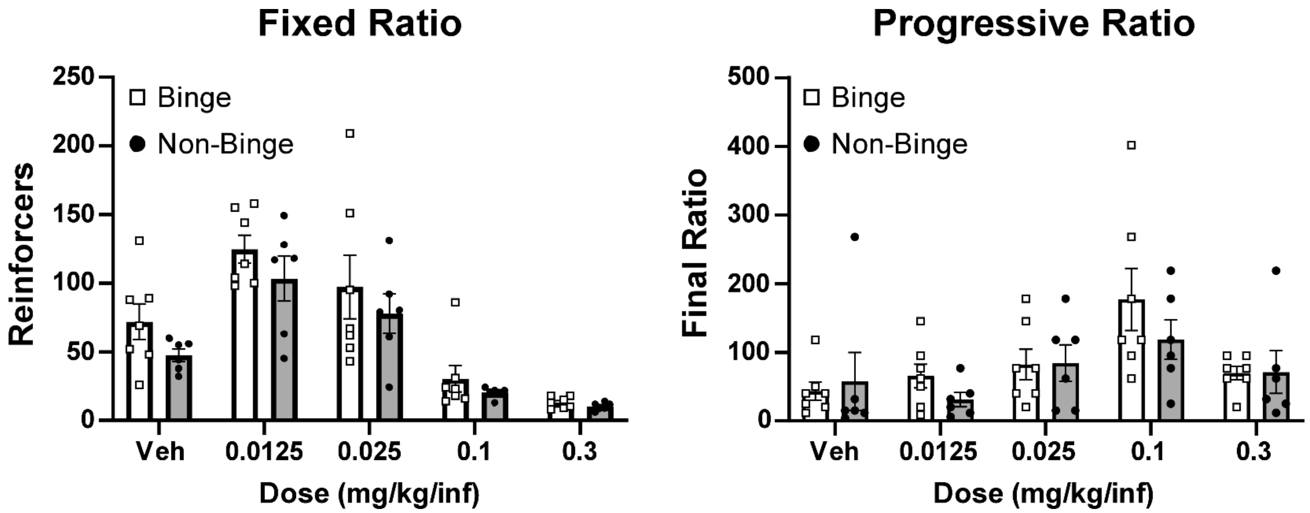


**Figure S4:** *Mean (±SEM) and individual reinforcers acquired in the Fixed Ratio dose substitution and on the final completed ratio in the Progressive Ratio dose substitution for the female and male rats trained on alpha-PVP who either met Binge criteria or did not (Non-Binge).*

### Impact of female odor on male IVSA:

There was no significant effect of session type (presence/absence of females in the operant chambers prior to running) on reinforcers acquired by male animals (**Figure S5**), although there was a significant effect of drug training Group confirmed [F (1, 13) = 11.17; P<0.01]. The posthoc test confirmed significant differences between drug training Groups for each session.

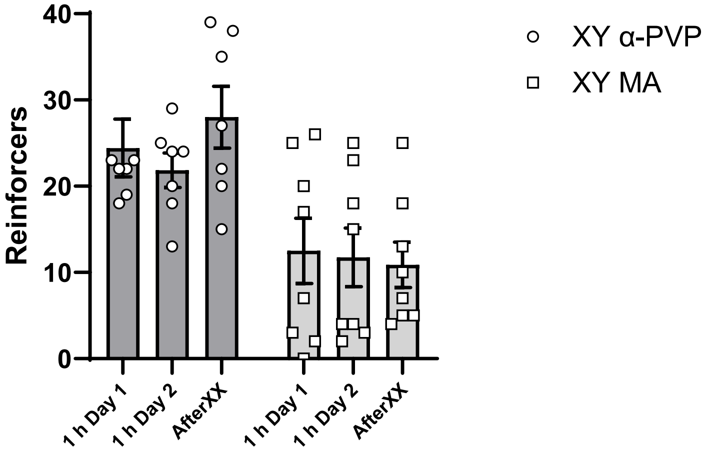


***Figure S5:*** *Mean (±SEM) infusions of their training drug acquired under a FR1 schedule of reinforcement in 1 h sessions by male rats originally trained on MA (N=8) or PVP (N=7). Sessions were run under baseline conditions, following the prior presence of female rats in the chambers or with no cleaning or bedding change prior to sessions.*

### α-pyrrolidinopropiophenone (α-PPP) and α-pyrrolidinohexiophenone (α-PHP):

*Fixed Ratio Dose Substitution*

As indicated in the main report there was no significant impact of the Sex factor (**Figure 4**) on responding for α-PPP and α-PHP across the tested doses (0.0125, 0.0250, 0.050 0.100, 0.300 mg/kg/infusion), however the dose-effect curves are presented in overlay here to emphasize the similarity (**Figure S6A**). Furthermore, although too few animals remained from the female MA group (N=3) for analysis the data are presented by training group in **Figure S6B** for completeness. In this analysis, there were also (N=5) female PVP, (N=7) male MA, and (N=7) male PVP rats remaining. The analysis of the male data (**Figure S6C**) confirmed that the male rats originally trained on MA obtained fewer reinforcers than the α-PVP trained males [Significant Effect of Dose $F(4, 48) = 11.21$; $P<0.0001$, of Training Drug $F(1, 47) = 5.075$; $P<0.05$, of Drug Identity: $F(1, 12) = 23.12$; $P<0.0005$ as well as of the interactions of Drug Identity and Dose: $F(4, 47) = 16.48$; $P<0.0001$, of Training Drug and Drug Identity: $F(1, 47) = 6.28$; $P<0.05$, and all three factors $F(4, 47) = 2.90$; $P<0.05$]. The post-hoc test of all possible comparisons did not confirm any significant differences between training groups at any dose of either drug. To further explore the main effects a two factor analysis treating training group and drug identity as a single group factor was conducted [Significant Effect of Dose $F(4, 95) = 14.71$; $P<0.0001$, of Group: $F(3, 24) = 4.84$; $P<0.01$ as well as of the interactions of Group and Dose: $F(12, 95) = 5.122$; $P<0.0001$]. The corresponding post-hoc test limited to orthogonal comparisons confirmed significant differences between Training Drug groups self-administering α-PHP at the 0.0125 mg/kg/infusion dose and when self-administering α-PPP at 0.025, 0.05 mg/kg/infusion doses. The MA-trained group self-administered significantly different numbers of infusions of α-PPP (0.05 vs 0.3 mg/kg/infusion) and α-PHP (0.0125 vs 0.3 mg/kg/infusion). The α-PHP-trained group self-administered significantly different numbers of

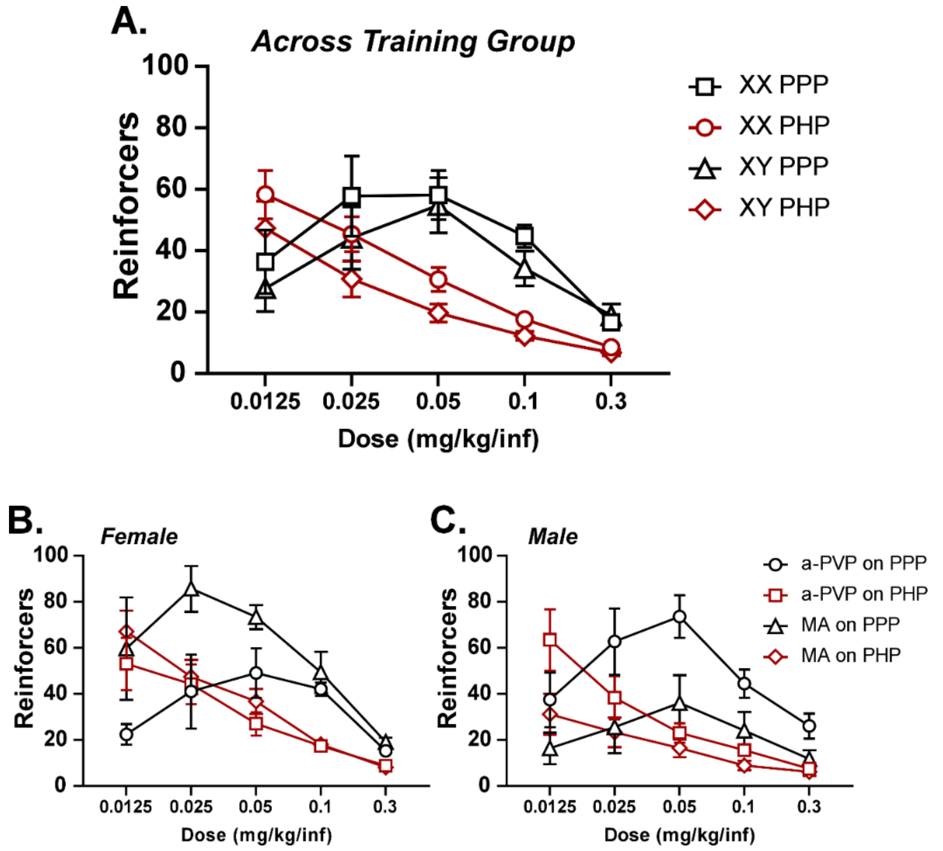


***Figure S6:*** *Mean (±SEM) infusions of α-PHP or α-PPP acquired under a FR1 schedule of reinforcement in 1 h sessions by **A)** female rats originally trained on **B)** MA (N=3) or PVP (N=5) and A) male rats originally trained on **C)** MA (N=8) or PVP (N=7).*

infusions of α-PPP (0.0125 vs 0.025, 0.05 mg/kg/infusion; 0.025 vs 0.3; 0.05 vs 0.1, 0.3 mg/kg) and α-PHP (0.0125 vs all other doses; 0.025 vs 0.3 mg/kg/infusion).

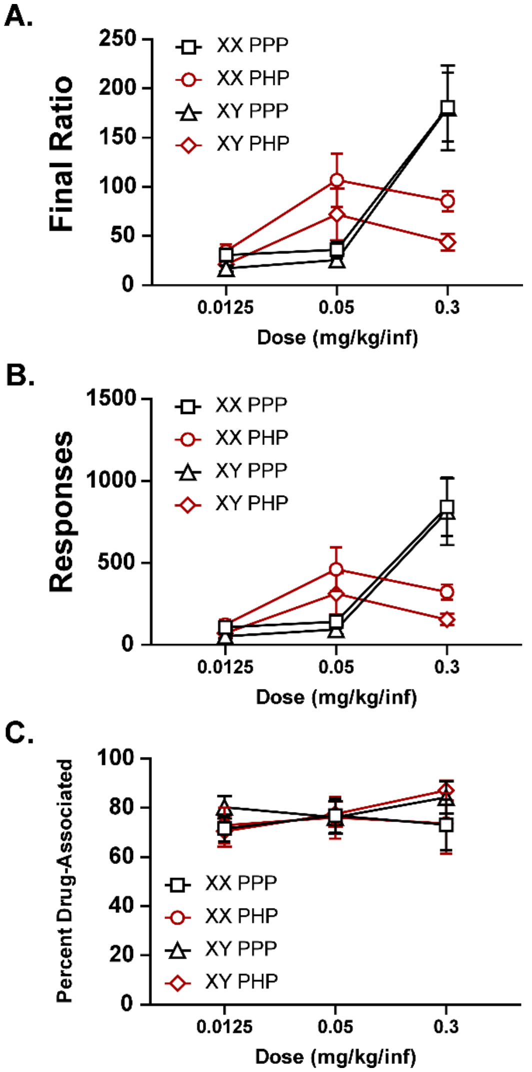


***Figure S7:*** *Mean (±SEM)* ***A)*** *final completed ratios,* ***B)*** *total drug-associated responses and* ***C)*** *percent drug-associated responding when either α-PHP or α-PPP was available under a PR schedule of reinforcement in 3 h sessions by male rats originally trained on MA (N=8) or PVP (N=7) and female rats originally trained on MA (N=3) or PVP (N=5).*

*Progressive Ratio*

The breakpoint (final completed ratio) reached in the PR study differed by dose and compound with the highest breakpoints achieved with the 0.3 mg/kg/infusion for α-PPP and the 0.05 mg/kg/infusion for α-PHP in each sex (**Figure S7A**). This further supports the conclusion from Figure S6A that α-PPP is less potent than α-PHP. A similar pattern held for total correct responses (**Figure S7B**) and the percentage of drug-associated responses did not vary by sex or drug (**Figure S7C**). As indicated in the main report, one additional female (α-PVP-trained group) lost patency during this study, thus N=7 for this analysis.

### Supplemental Discussion

Roughly half of individuals trained in the IVSA of α-PVP (female rats) or the closely related MDPV compound (male rats) exhibited a binge-like acquisition day in our prior studies. This appeared as a day of high intake, after which lower amounts are self-administered from session to session in a relatively stable pattern. In one additional study, an increase in the female group for the third day of acquisition of IVSA suggests a similar binge-like acquisition pattern, although the authors did not explicate the trend with individual data (Marusich et al., 2021).The ratio of binge-like acquisition patterns in prior work was 15/25 in female Wistar rats self-administering alpha-PVP (0.05 mg/kg/infusion) (Javadi-Paydar et al., 2018) and in 13/26 male Wistar rats self-administering MDPV (0.05 mg/kg/infusion) (Aarde et al., 2015). The ratio observed here overall (8/15) is quite similar and given the group sizes, there is no evidence of any sex difference. Overall, there was no evidence for any lasting impact of the binge phenotype, i.e. in the FR1 dose-substitution (as found previously for female rats (Javadi-Paydar et al., 2018)) or in post-binge training days (as was found for males (Aarde et al., 2015)). These results confirm that our prior results were not an artifact of larger operant chambers which included a running wheel which was used in those prior studies.

The lack of any difference in male rat's intravenous self-administration of either methamphetamine or α-PVP did not support the hypothesis that being run in operant boxes after female animals alters this behavior. This is an initial minor attempt to address cautions long discussed, and recently published (Dalla et al., 2024), in the context of the Sex As a Biological Variable mandate of the NIH. There are, however, numerous caveats. This was conducted post-acquisition and when patterns had stabilized, it may therefore not apply to initial acquisition. The sample was relatively small for each drug and 3/7 of the α-PVP rats obtained more infusions after females were run than any rat in the comparison sessions.

### Supplemental Literature Cited


Aarde, S.M., Huang, P.K., Dickerson, T.J., Taffe, M.A., 2015. Binge-like acquisition of 3,4-methylenedioxypyrovalerone (MDPV) self-administration and wheel activity in rats. Psychopharmacology 232(11), 1867-1877, doi: 10.1007/s00213-014-3819-4.

Javadi-Paydar, M., Harvey, E.L., Grant, Y., Vandewater, S.A., Creehan, K.M., Nguyen, J.D., Dickerson, T.J., Taffe, M.A., 2018. Binge-like acquisition of alpha-pyrrolidinopentiophenone (alpha-PVP) self-administration in female rats. Psychopharmacology 235(8), 2447-2457, doi: 10.1007/s00213-018-4943-3.

Marusich, J.A., Gay, E.A., Watson, S.L., Blough, B.E., 2021. Alpha-pyrrolidinopentiophenone and mephedrone self-administration produce differential neurochemical changes following short- or long-access conditions in rats. Eur J Pharmacol 897, 173935, doi: 10.1016/j.ejphar.2021.173935.
